# The decision of male medaka to mate or fight depends on two complementary androgen signaling pathways

**DOI:** 10.1101/2024.01.10.574747

**Authors:** Yuji Nishiike, Kataaki Okubo

## Abstract

Adult male animals typically court and attempt to mate with females, while attacking other males. Emerging evidence from mice indicates that neurons expressing the estrogen receptor ESR1 in behaviorally relevant brain regions play a central role in mediating these mutually exclusive behavioral responses to conspecifics. However, the findings in mice are unlikely to apply to most other vertebrates, where androgens — rather than estrogens — have been implicated in male behaviors. Here we report that male medaka (*Oryzias latipes*) lacking one of the two androgen receptor subtypes (Ara) are less aggressive toward other males and instead actively court them, while those lacking the other subtype (Arb) are less motivated to mate with females and conversely attack them. These findings indicate that, in male medaka, the Ara- and Arb-mediated androgen signaling pathways facilitate appropriate behavioral responses, while simultaneously suppressing inappropriate responses, to males and females, respectively. Notably, males lacking either receptor retain the ability to discriminate the sex of conspecifics, suggesting a defect in the subsequent decision-making process to mate or fight. We further show that Ara and Arb are expressed in intermingled but largely distinct populations of neurons, and stimulate the expression of different behaviorally relevant genes including galanin and vasotocin, respectively. Collectively, our results demonstrate that male teleosts make adaptive decisions to mate or fight as a result of the activation of one of two complementary androgen signaling pathways, depending on the sex of the conspecific that they encounter.

## Introduction

Across species, adult males typically court and attempt to mate with females, while directing aggression toward other males. The question of how these mutually exclusive behavioral responses to conspecifics are generated has long been a topic of great interest (1, 2). The underlying neural mechanisms, while extensively studied in flies (3, 4), have generally remained elusive in vertebrates. Recent studies in mice, however, have yielded important findings: neurons expressing the estrogen receptor (ESR) subtype ESR1 in the medial preoptic area (MPOA) and the ventrolateral part of the ventromedial hypothalamus (VMHvl), which evoke mating and aggression, respectively, inhibit each other to ensure mutually exclusive displays of these behaviors (5, 6).

Nevertheless, the specific role of estrogen/ESR1 signaling in these neurons is still unclear. It has been consistently shown in rodents that estrogen/ESR1 signaling is essential for male behaviors (7–9). Specifically, testosterone secreted from the fetal testis is converted to estradiol-17β (E2) in the developing brain, which then acts through ESR1 to organize the neural substrates that later mediate male-typical behaviors. In addition, adult testicular testosterone, after conversion to E2, activates the neural substrates through ESR1 to achieve male-typical behaviors (1, 9–12). Despite these well-established findings, the neural mechanisms by which estrogen/ESR1 signaling mediates male behaviors, including its target genes, are largely undefined (12).

More importantly, the neural circuits that mediate social behaviors, including mating and aggression (often referred to as the “social behavior network”), seem to be highly conserved across vertebrates (13, 14); in many non-rodent species including humans, most birds, and teleost fish, however, androgen/androgen receptor (AR) signaling — rather than estrogen/ESR signaling — has been implicated in male behaviors (i.e., testicular androgens act directly through AR without conversion to estrogens) (15–18). This suggests that the neural underpinnings of male-typical behavioral responses observed in mice do not apply to these species.

In teleosts, 11-ketotestosterone (11KT), which cannot be converted to estrogen, is the primary testicular androgen, and treating females with 11KT as adults effectively induces male-typical behaviors, including courtship and aggression (18–21). These facts suggest that both organization and activation of the neural substrates for male behaviors rely solely on androgen/AR signaling in adulthood. As such, teleosts provide simple and easy-to-manipulate model systems for studying the neural and hormonal mechanisms underlying male-typical behavioral responses. Most teleost species have two AR subtypes, Ara and Arb, resulting from a whole-genome duplication that occurred early in the teleost lineage (22, 23). The role of each AR in testicular development and secondary sexual characteristics has recently been studied in AR-deficient models of cichlid (*Astatotilapia burtoni*) and medaka (*Oryzias latipes*) (24, 25). In the present study, we have studied the role of androgen/AR signaling in male-typical behavioral responses by investigating the behavior of AR-deficient male medaka toward conspecifics. We find, to our surprise, that males deficient in Ara frequently court other males, while males deficient in Arb attack females. Our findings provide evidence that two functionally distinct AR signaling pathways act in a complementary manner in males to generate appropriate behavioral responses during social encounters.

## Results

### Androgen/Ara signaling facilitates male aggression toward other males

We generated medaka deficient in *ara* and *arb* using clustered regularly interspaced short palindromic repeats (CRISPR)/CRISPR-associated protein 9 (Cas9) genome editing. To ensure the reproducibility of the observed behavioral phenotypes, two independent medaka lines were established for *ara* (Δ326 and Δ325; Fig. S1) and for *arb* (Δ10 and Δ11; Fig. S2) and used for subsequent analyses. Note that this paper follows the nomenclature of teleost ARs based on their orthology/paralogy relationships (23, 25), and the designations Ara and Arb are opposite to those in our earlier publications (e.g., 19, 26–29).

We first used these medaka to investigate the consequences of impaired Ara- and Arb-mediated androgen signaling on male aggression toward other males. The aggressive behavior of teleosts, including medaka, involves five types of behavioral act: chases, fin displays, circles, strikes, and bites (Fig. 1A) (19, 30). Tests of aggressive behavior among grouped *ara*-deficient (*ara*^−/−^) males revealed that they engaged in all of these aggressive acts less frequently than their wild-type (*ara*^+/+^) siblings (although not significantly for some acts) in both the Δ326 and Δ325 lines (Fig. 1 B–D). In contrast, similar behavioral testing in *arb*-deficient (*arb*^−/−^) males showed no significant difference in the frequency of any aggressive acts between these males and their wild-type (*arb*^+/+^) siblings in both the Δ10 and Δ11 lines (Fig. 1 B, E, and F). Taken together, these results indicate that male aggression toward other males is primarily facilitated by androgen/Ara signaling.

**Fig. 1.**
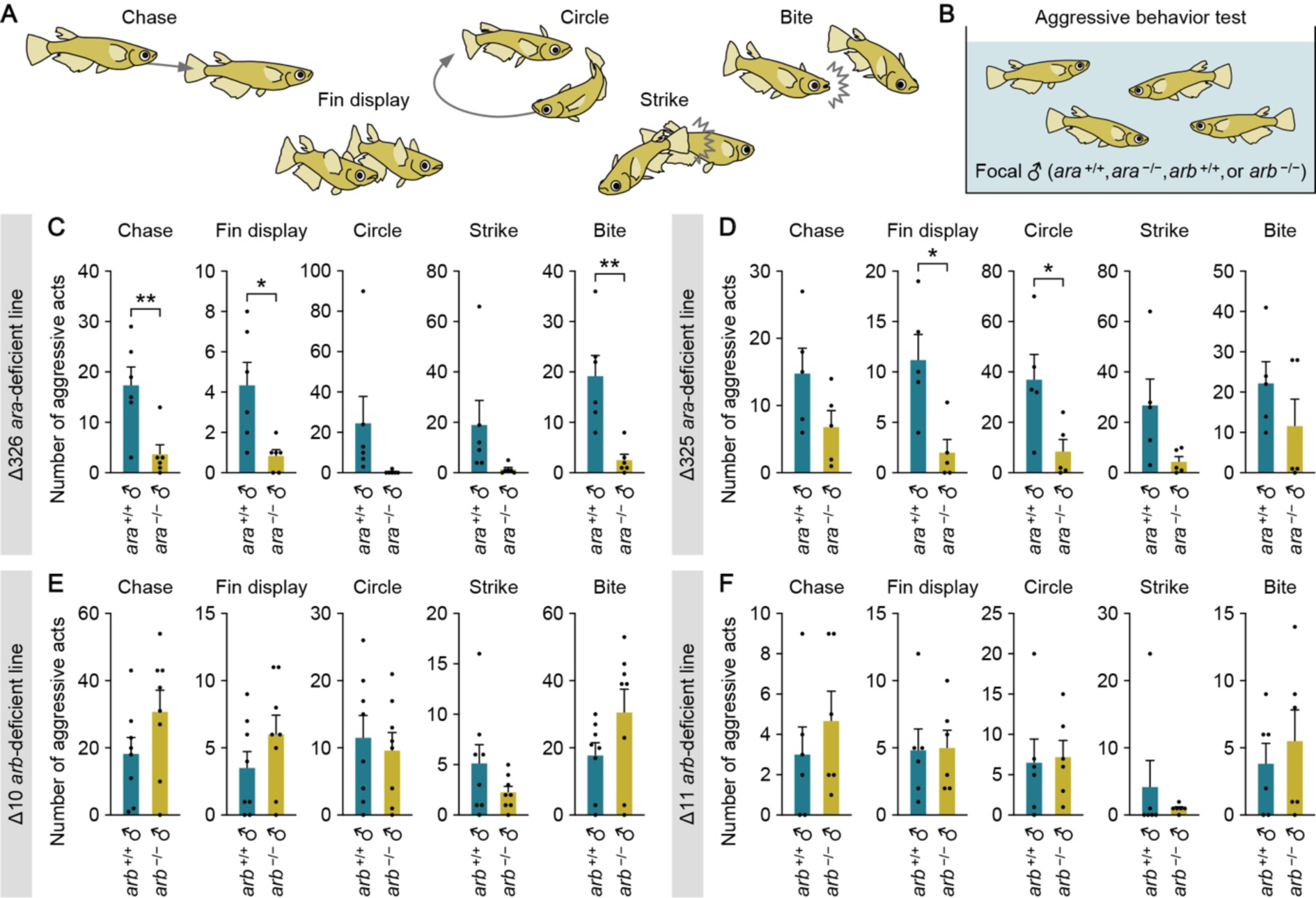
Androgen/Ara signaling facilitates male aggression toward other males. (**A**) Schematic of the five types of aggressive act. (**B**) Set-up for testing aggressive behavior among grouped males. (**C** and **D**) Total number of each aggressive act observed among *ara*^+/+^ males and among *ara*^−/−^ males in the tank. Results from both Δ326 (n = 6 per genotype; C) and Δ325 (n = 5 per genotype; D) *ara*-deficient lines are shown. (**E** and **F**) Total number of each aggressive act observed among *arb*^+/+^ males and among *arb*^−/−^ males in the tank. Results from both Δ10 (n = 8 per genotype; E) and Δ11 (n = 6 per genotype; F) *arb*-deficient lines are shown. Statistical differences were calculated by unpaired *t* test, with Welch’s correction where appropriate (C–F). Error bars represent SEM. \**P* < 0.05, \*\**P* < 0.01.

### Androgen/Arb signaling facilitates male mating with females

Next, we assessed the impact of impaired Ara- and Arb-mediated androgen signaling on male mating with females. The mating behavior of medaka follows a stereotyped pattern, wherein a series of followings, courtship displays, and wrappings by the male leads to spawning (31, 32) (Fig. 2A). Tests of the mating behavior of *ara*^−/−^ males paired with a stimulus female showed that they initiated followings, courtship displays, and wrappings with latencies comparable to those of *ara*^+/+^ males in both the Δ326 and Δ325 lines (Fig. 2 B–D). In addition, the majority (>80%) of *ara*^−/−^ males spawned during the test period (Fig. 2 C and D). These observations indicate that loss of androgen/Ara signaling does not affect male mating with females.

**Fig. 2.**
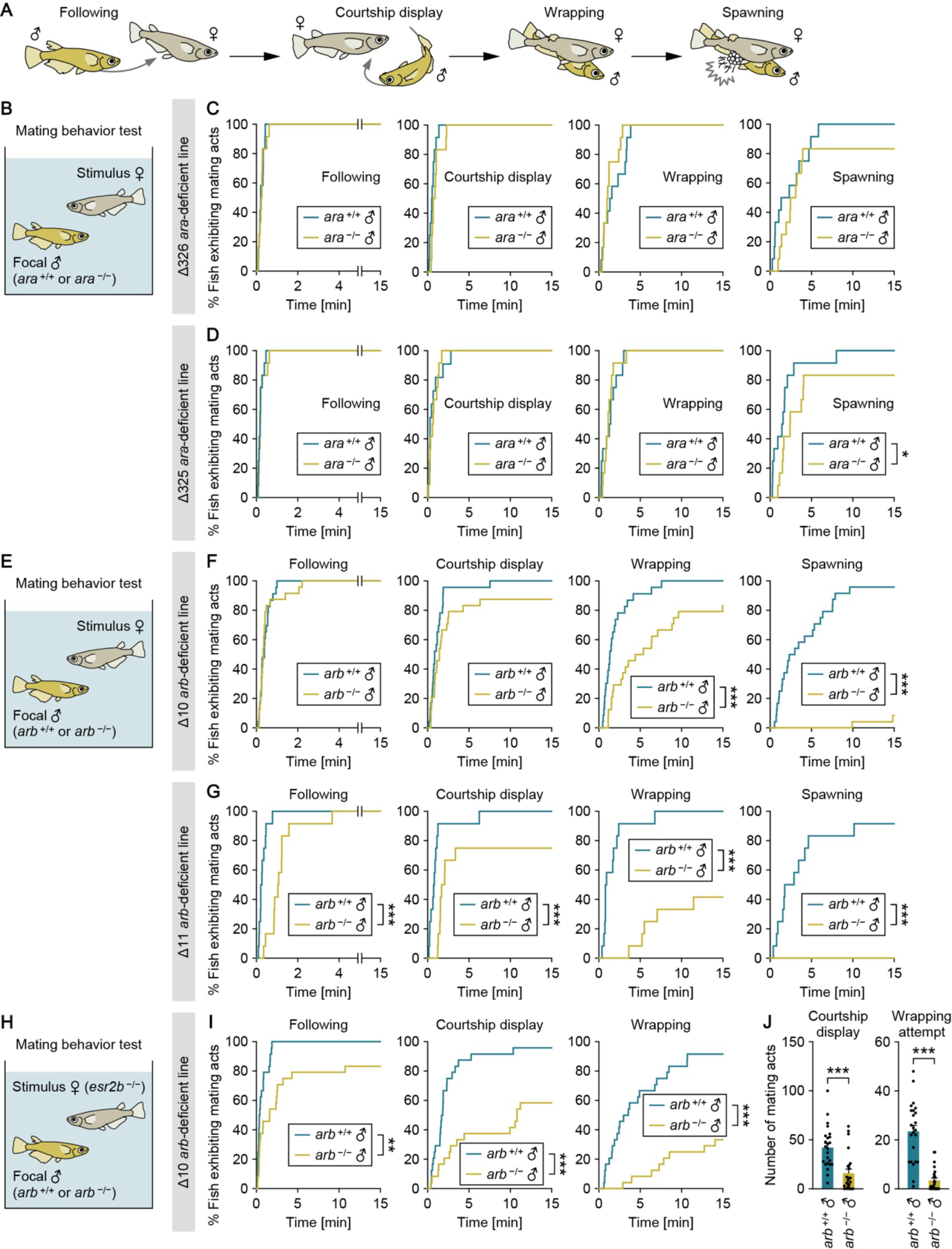
Androgen/Arb signaling facilitates male mating with females. (**A**) Schematic of a sequence of male-typical mating acts. (**B**) Set-up for testing the mating behavior of *ara*^+/+^ and *ara*^−/−^ males. (**C** and **D**) Latency of *ara*^+/+^ and *ara*^−/−^ males to initiate each mating act toward the stimulus female. Results from both Δ326 (n = 12 per genotype; C) and Δ325 (n = 12 per genotype; D) *ara*-deficient lines are shown. (**E**) Set-up for testing the mating behavior of *arb*^+/+^ and *arb*^−/−^ males. (**F** and **G**) Latency of *arb*^+/+^ and *arb*^−/−^ males to initiate each mating act toward the stimulus female. Results from both Δ10 (n = 23, and 24 for *arb*^+/+^ and *arb*^−/−^, respectively; F) and Δ11 (n = 12 for each genotype; G) *arb*-deficient lines are shown. (**H**) Set-up for the additional mating behavior test using an *esr2b*-deficient female as the stimulus. (**I**) Latency of the focal *arb*^+/+^ and *arb*^−/−^ males (Δ10 line; n = 24 per genotype) to initiate each mating act. (**J**) Number of each mating act performed. Statistical differences were calculated by Gehan-Breslow-Wilcoxon test (C, D, F, G, and I) and unpaired *t* test (J). Error bars represent SEM. \**P* < 0.05, \*\**P* < 0.01, \*\*\**P* < 0.001.

Conversely, behavioral testing in *arb*^−/−^ males of the Δ10 line paired with a stimulus female revealed significantly longer latencies to initiate wrappings and spawn, with less than 10% of them spawning successfully (Fig. 2 E and F). Similar results were obtained in the Δ11 line, where *arb*^−/−^ males showed significantly longer latencies to followings and courtship displays, in addition to wrappings and spawning, and none of them spawned in the test period (Fig. 2 E and G). These observations suggest that loss of androgen/Arb signaling renders males less motivated to mate with females.

However, given that *arb*^−/−^ males lack male-typical secondary sexual characteristics in fin morphology (25), we considered that their mating defects might be the result of females being less sexually attracted to *arb*^−/−^ males. To address this possibility, their mating behavior was tested again using females deficient in the ESR subtype Esr2b, which are completely unreceptive to male courtship (20), as stimulus females. This test, which enabled us to assess the motivation of males to mate with females without the influence of female receptivity, revealed that *arb*^−/−^ males were indeed less motivated to mate with females. Specifically, *arb*^−/−^ males showed significantly longer latencies to initiate followings, courtship displays, and wrappings, and significantly fewer courtship displays and wrapping attempts than *ara*^+/+^ males in both the Δ10 (Fig. 2 H–J) and Δ11 lines (Fig. S3 A–C). Similar tests on *ara*^−/−^ males showed that they did not differ significantly from *ara*^+/+^ males in any measure (Fig. S3 D–H). Collectively, these findings indicate that the motivation of males to mate with females is primarily facilitated by androgen/Arb signaling.

### Adult androgen/AR signaling elicits male-typical behavioral responses even in females

Female teleosts, when treated with 11KT as adults, display male-typical aggressive and mating behavior while retaining their ovaries (18–21, 33). We considered that, if the above conclusions apply to females as well as males, the male-typical aggressive and mating behaviors observed in 11KT-treated females are likely to be elicited through Ara and Arb, respectively. To address this possibility and further verify our preceding findings, we investigated whether *ara*^−/−^ (Δ326 line) and *arb*^−/−^ (Δ10 line) females treated with 11KT as adults exhibit male-typical aggressive and mating behaviors.

In tests of aggressive behavior towards a stimulus male, none of the 11KT-treated *ara*^−/−^ females exhibited any aggressive acts, whereas more than 60% of the control 11KT-treated *ara*^+/+^ females did (Fig. S4 A and B); in addition, there were significant differences between the two genotypes in the frequency of fin displays and bites (Fig. S4C). In contrast, more than 50% of 11KT-treated *arb*^−/−^ females exhibited aggressive acts towards a stimulus male, and these females did not differ significantly from the control *arb*^+/+^ females in the frequency of any acts (Fig. S4 D–F). We further examined aggressive behavior among grouped females treated with 11KT, but hardly any aggressive acts were observed regardless of strain or genotype (Fig. S4 G–I), probably because these females were not fully masculinized in appearance and did not recognize each other as targets for attack.

In tests of mating behavior toward a stimulus female, 11KT-treated *ara*^−/−^ females initiated followings, courtship displays, and wrappings with latencies comparable to those of *ara*^+/+^ females (Fig. S4 J and K). In contrast, only a fraction of 11KT-treated *arb*^−/−^ females performed followings or courtship displays to the stimulus female, and their latencies to these acts were significantly longer than those of *arb*^+/+^ females (Fig. S4 L and M). Furthermore, none of the *arb*^−/−^ females engaged in wrapping (Fig. S4M). Together, these results suggest that Ara- and Arb-mediated androgen signaling in adulthood elicit male-typical patterns of aggression and mating, respectively, even in females (i.e., regardless of genetic or phenotypic sex).

### *ara*-deficient males frequently attempt to mate with other males

Curiously, in examining the aggressive behavior of *ara*^−/−^ males, we noted that they frequently exhibited mating acts to other males. To explore this observation further, we tested and quantified the mating behavior of *ara*^−/−^ males paired with a stimulus male (Fig. 3 A and B). The results showed that *ara*^−/−^ males performed courtship displays and attempted wrappings to the stimulus male more frequently and had a shorter latency to wrapping attempts as compared with their *ara*^+/+^ siblings (although not significantly for some results) in both the Δ326 and Δ325 lines (Fig. 3 C–F). These data collectively indicate that *ara*^−/−^ males not only are less aggressive toward other males but also frequently attempt to mate with them.

**Fig. 3.**
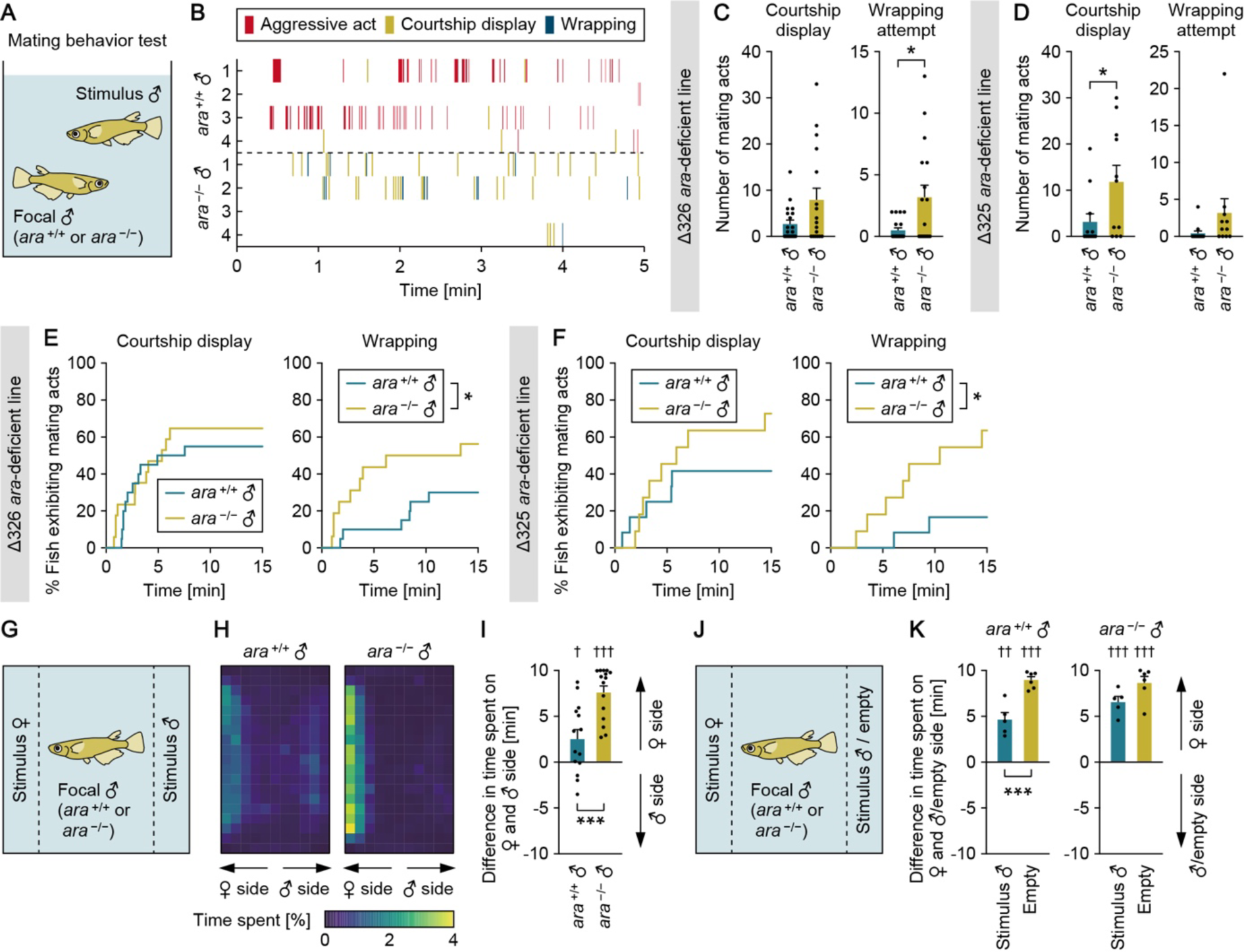
*ara*-deficient males frequently attempt to mate with other males. (**A**) Set-up for testing the mating behavior of *ara*^+/+^ and *ara*^−/−^ males toward other males. (**B**) Raster plots of behavioral responses of males to the stimulus male. Data from four representative males of each genotype (*ara*^+/+^ and *ara*^−/−^; Δ326 line) are shown. (**C** and **D**) Number of each mating act performed by *ara*^+/+^ and *ara*^−/−^ males toward the stimulus male. Results from both Δ326 (n = 20 and 17 for *ara*^+/+^ and *ara*^−/−^, respectively; C) and Δ325 (n = 12 and 11 for *ara*^+/+^ and *ara*^−/−^, respectively; D) *ara*-deficient lines are shown. (**E** and **F**) Latency of males to initiate each mating act. Results from both Δ326 (E) and Δ325 (F) lines are shown. (**G**) Set-up for the three-chamber test to assess the sex discrimination ability of *ara*^+/+^ and *ara*^−/−^ males. (**H**) Heat maps depicting the time spent by the focal males (Δ326 line; n = 14 and 15 for *ara*^+/+^ and *ara*^−/−^, respectively) in each location of the test chamber. (**I**) Difference in time spent by the focal males on the stimulus female versus the stimulus male side. (**J**) Set-up for the additional three-chamber test to assess the influence of removing the stimulus male. (**K**) Difference in time spent by the focal *ara*^+/+^ and *ara*^−/−^ males (Δ326 line) on the stimulus female versus the stimulus male (n = 5) or empty (n = 6) side. Statistical differences were calculated by unpaired *t* test, with Welch’s correction where appropriate (C, D, and comparisons between genotypes in I and K), Gehan-Breslow-Wilcoxon test (E and F), and one-sample *t* test (comparisons against the null hypothesis of no difference in I and K). Error bars represent SEM. \**P* < 0.05, \*\**P* < 0.01, \*\*\**P* < 0.001 between genotypes. ^†^*P* < 0.05, ^††^*P* < 0.01, ^†††^*P* < 0.001 against the null hypothesis.

What behavioral mechanisms, then, underlie the male-directed mating attempts of *ara*^−/−^ males? We considered that there are three possible explanations: 1) *ara*^−/−^ males cannot discriminate the sex of conspecifics and consequently misidentify males as females; 2) they can discriminate sex, but cannot make appropriate decisions about how to respond to male conspecifics; and 3) they are excessively sexually aroused and seek to mate with conspecifics regardless of their sex. As this last possibility seemed unlikely because *ara*^−/−^ males were not overly motivated to mate with females, we assessed the first and second possibilities by investigating the sex discrimination ability of *ara*^−/−^ males in a three-chamber test with a stimulus female in one side chamber and a stimulus male in the other (Fig. 3G). Similar to *ara*^+/+^ males, *ara*^−/−^ males spent significantly more time near the stimulus female than near the male (Fig. 3 H and I), suggesting that *ara*^−/−^ males can discriminate the sex of conspecifics.

Unexpectedly, *ara*^−/−^ males spent even more time near the female than did *ara*^+/+^ males (Fig. 3 H and I). This observation could be explained by assuming that *ara*^−/−^ males either have an increased preference for females or avoid contact with other males. Given that *ara*^−/−^ males were not more motivated to mate with females and were less aggressive toward other males, their decreased propensity to attack other males probably led them to spend relatively more time near the female. We tested this presumption by an additional three-chamber test in which the stimulus male was removed to leave an empty side chamber. Whereas *ara*^+/+^ males spent significantly more time near the female in the absence than in the presence of the stimulus male, *ara*^−/−^ males showed no difference in their behavior with or without the stimulus male (Fig. 3 J and K), again suggesting a reduced propensity of *ara*^−/−^ males to attack other males. In summary, *ara*^−/−^ males actively court and attempt to mate with other males even though they retain the ability to discriminate the sex of conspecifics. Thus, androgen/Ara signaling presumably prevents the maladaptive decision to engage in mating with other males.

We also investigated the mating behavior of *arb*^−/−^ males toward a stimulus male. In contrast to *ara*^−/−^ males, *arb*^−/−^ males performed courtship displays and attempted wrappings less frequently and with longer latencies as compared with *arb*^+/+^ males (Fig. S5). Therefore, *arb*^−/−^ males are less inclined than *arb*^+/+^ males to mate with males or females, suggesting that mating behavior toward females and males is stimulated by common neural substrates involving androgen/Arb signaling.

### *arb*-deficient males attack females

In examining the mating behavior of *arb*^−/−^ males, we also noted that they attacked females (in general, male medaka rarely attack females). To explore this observation, we tested and quantified the aggressive behavior of *arb*^−/−^ males toward a stimulus female (Fig. 4 A and B). While none of the *arb*^+/+^ males exhibited aggressive acts toward the stimulus female, more than half of the *arb*^−/−^ males from both the Δ10 and Δ11 lines did (Fig. 4 C and D). Furthermore, they exhibited all five types of aggressive act rather than specific acts, with significant increases in chases, fin displays, circles, and bites in the Δ10 line, and in chases and bites in the Δ11 line (Fig. 4 E and F). These results demonstrate that *arb*^−/−^ males not only are less motivated to mate with females, but also are aggressive toward them. *ara*^−/−^ males, on the other hand, did not show any aggressive acts toward the stimulus female (Fig. S6 A– C).

**Fig. 4.**
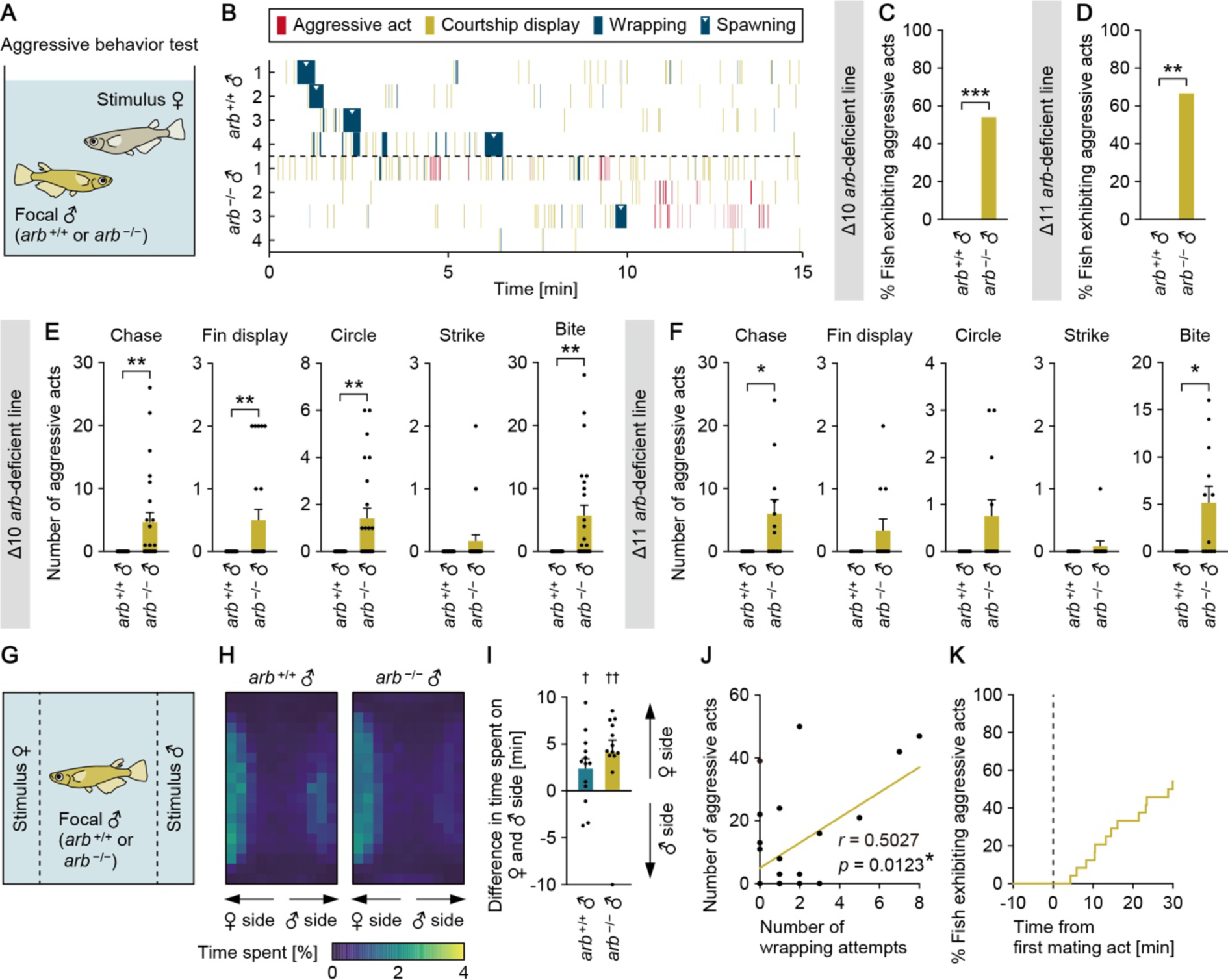
*arb*-deficient males attack females. (**A**) Set-up for testing the aggressive behavior of *arb*^+/+^ and *arb*^−/−^ males toward females. (**B**) Raster plots of behavioral responses of males to the stimulus female. Data from four representative males of each genotype (*arb*^+/+^ and *arb*^−/−^; Δ10 line) are shown. (**C** and **D**) Percentage of *arb*^+/+^ and *arb*^−/−^ males exhibiting any aggressive act toward the stimulus female. Results from both Δ10 (n = 23 and 24 for *arb*^+/+^ and *arb*^−/−^, respectively; C) and Δ11 (n = 12 for each genotype; D) *arb*-deficient lines are shown. (**E** and **F**) Number of each aggressive act performed. Results from both Δ10 (E) and Δ11 (F) lines are shown. (**G**) Set-up for the three-chamber test to assess the sex discrimination ability of *arb*^+/+^ and *arb*^−/−^ males. (**H**) Heat maps depicting the time spent by the focal males (Δ10 line; n = 13 per genotype) in each location of the test chamber. (**I**) Difference in time spent by the focal males on the stimulus female versus the stimulus male side. (**J**) Scatter plot of the number of aggressive acts and wrapping attempts performed by *arb*^−/−^ males (Δ10 line; n = 24) toward the stimulus female. Each dot represents one focal male; the unbroken line represents the regression line. (**K**) Latency from the first mating act to the initiation of aggressive acts. Statistical differences were calculated by Fisher’s exact test (C and D), unpaired *t* test, with Welch’s correction where appropriate (E, F, and comparisons between genotypes in I), one-sample *t* test (comparisons against the null hypothesis of no difference in I), and pairwise Pearson correlation (J). Error bars represent SEM. \**P* < 0.05, \*\**P* < 0.01, \*\*\**P* < 0.001 between genotypes. ^†^*P* < 0.05, ^††^*P* < 0.01 against the null hypothesis.

This finding prompted us to investigate the sex discrimination ability of *arb*^−/−^ males, as we did for *ara*^−/−^ males. In the three-chamber test with a stimulus male in one side chamber and a stimulus female in the other, *arb*^−/−^ males, as well as *arb*^+/+^ males, spent more time near the stimulus female; in addition, the degree of time spent near the female was comparable between *arb*^+/+^ and *arb*^−/−^ males (Fig. 4 G–I). Thus, *arb*^−/−^ males attack females despite being able to discriminate the sex of conspecifics, suggesting that androgen/Arb signaling prevents the maladaptive decision to attack females.

We further noticed that the individual *arb*^−/−^ males that engaged more frequently in mating acts toward the stimulus females also showed more frequent aggressive acts toward them (e.g., compare the *arb*^−/−^ males in rows 1–3 versus row 4 in Fig. 4B). Analysis of the correlation between the frequency of aggressive acts and wrapping attempts indeed showed a positive correlation between them (Fig. 4J). In addition, none of the *arb*^−/−^ males initiated aggressive acts prior to mating acts (Fig. 4K and Fig. S6 D and E); when they spawned, however, all of them turned to aggressive acts immediately (within 3 min) (4/4 and 3/3 individuals in the Δ10 and Δ11 lines, respectively; e.g., see the *arb*^−/−^ male in row 3 in Fig. 4B). Coupled with the finding in mice and flies that interactions with females increase intrasexual aggression in males (34, 35), these observations led us to assume that *arb*^−/−^ males may be aggressive during the consummatory phase of mating. We tested this idea by assessing the aggression of *arb*^−/−^ males paired with *esr2b*-deficient females, which are unreceptive to male courtship and prevent males from proceeding to the consummatory phase (20). As expected, *arb*^−/−^ males showed scarcely any aggressive acts toward the *esr2b*-deficient females (Fig. S6 F–J), thereby supporting our assumption.

In summary, these results demonstrate that *arb*^−/−^ males are aggressively aroused upon mating with females, suggesting partially shared neural substrates for mating and aggression. Androgen/Arb signaling may act on these shared neural substrates to promote a state of sexual arousal, while simultaneously suppressing aggressive arousal, toward females.

### *ara* and *arb* are preferentially expressed in different neurons

Overall, the above findings indicate that androgen signaling via Ara and Arb mediates behavioral responses to male and female conspecifics, respectively. This raises the new question of how Ara and Arb might mediate such different behavioral responses, despite sharing common ligands, binding sites, and even expression sites in the medaka brain (26). To answer this question, we analyzed the coexpression of *ara* and *arb* at the cellular level by double in situ hybridization, which showed that these genes are indeed expressed together in many brain regions, but mainly in distinct populations of intermingled neurons. Neurons coexpressing *ara* and *arb* accounted for less than 30% of all neurons expressing these genes in any brain nucleus (28% and 23% in the PPa and NPT, respectively), and were rarely seen in the Vs/Vp (4.5%), PMp (2.8%), PPp (4.0%), and NVT (4.4%) (Fig. 5A and Fig. S7A; see Table S1 for abbreviations of brain nuclei). These data suggest that the Ara and Arb signaling pathways mediate different behavioral responses by acting independently of each other in different populations of neurons.

**Fig. 5.**
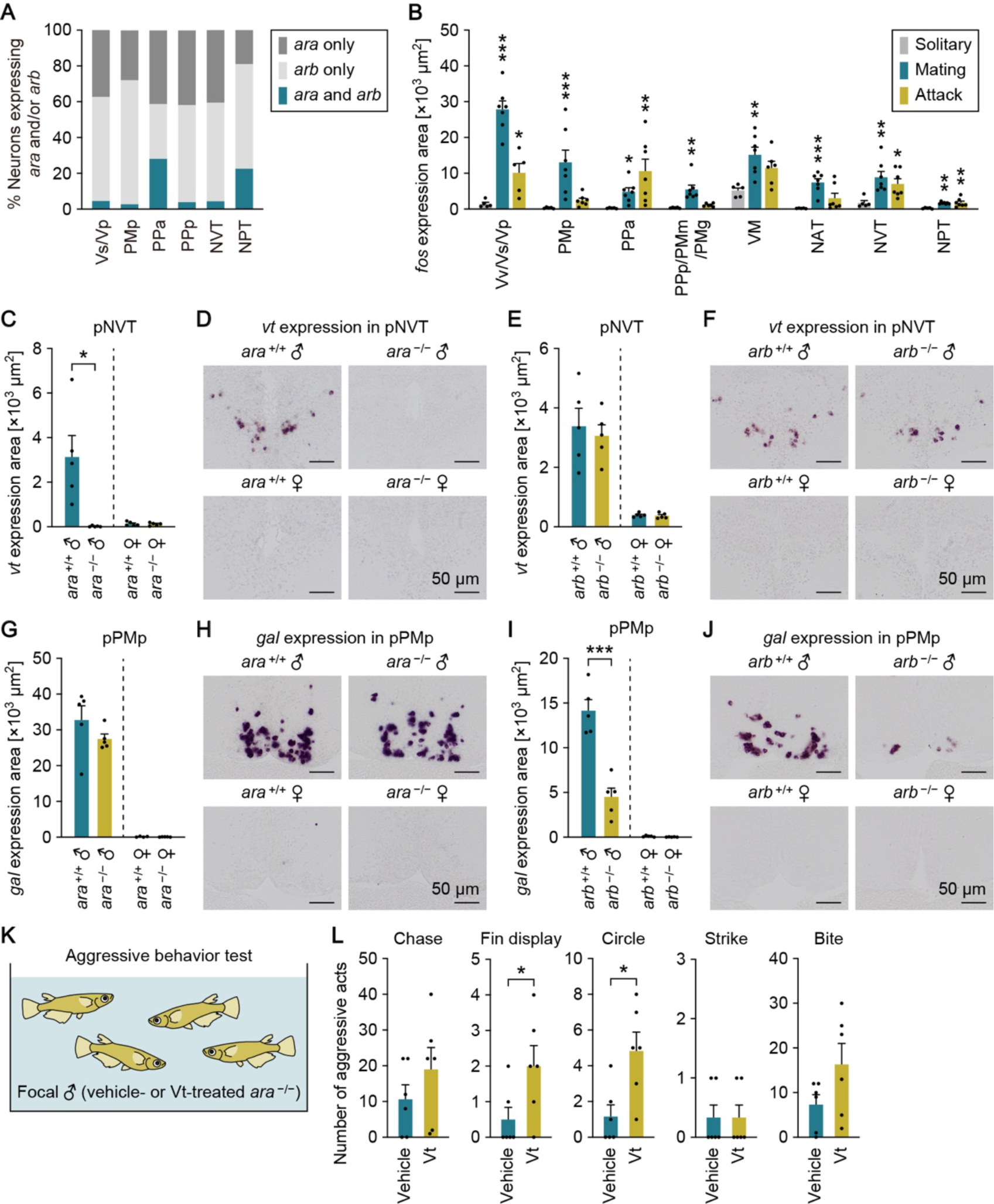
Ara and Arb are expressed in different neurons and stimulate different behaviorally relevant genes. (**A**) Percentage of neurons expressing *ara* only, *arb* only, or both in each brain nucleus of wild-type males. (**B**) Total area of *fos* expression signal in brain nuclei (where *ara* and/or *arb* are expressed) of wild-type solitary males (n = 5), males that mated with females (n = 7), and males that attacked other males (n = 7 except for Vv/Vs/Vp, PPp/PMm/PMg, and VM, where n = 5, 6, and 6, respectively). (**C** and **D**) Total area (C) and representative images (D) of *vt* expression signal in the pNVT of males and females of *ara*^+/+^ and *ara*^−/−^ fish (Δ326 line; n = 5 per sex per genotype). (**E** and **F**) Total area (E) and representative images (F) of *vt* expression signal in the pNVT of males and females of *arb*^+/+^ and *arb*^−/−^ fish (Δ10 line; n = 5 per sex per genotype). (**G** and **H**) Total area (G) and representative images (H) of *gal* expression signal in the pPMp of males and females of *ara*^+/+^ and *ara*^−/−^ fish (Δ326 line; n = 5 per sex per genotype, except n = 4 for *ara*^+/+^ females). (**I** and **J**) Total area (I) and representative images (J) of *gal* expression signal in the pPMp of males and females of *arb*^+/+^ and *arb*^−/−^ fish (Δ10 line; n = 5 per sex per genotype). (**K**) Set-up for testing aggressive behavior among *ara*^−/−^ males treated with vehicle alone or Vt peptide. (**L**) Total number of each aggressive act observed among *ara*^−/−^ males in the tank (Δ326 line; n = 6 per treatment). All scale bars are 50 μm. For abbreviations of brain nuclei, see Table S1. Statistical differences were calculated by Dunnet’s or Dunn’s post hoc test (B) and unpaired *t* test, with Welch’s correction where appropriate (C, E, G, I, and L). Error bars represent SEM. \**P* < 0.05, \*\**P* < 0.01, \*\*\**P* < 0.001.

This notion was further supported by the results of in situ hybridization analysis of *arb* expression in *ara*^−/−^ male brains and *ara* expression in *arb*^−/−^ male brains, which revealed that the expression of either gene in any brain nucleus did not differ significantly between these males and their wild-type siblings (Fig. S7 B and C). Loss of one AR subtype, therefore, does not lead to a likely functional compensation via upregulation of the other paralogous subtype, again suggesting the independence of the Ara and Arb signaling pathways.

### The two androgen/AR signaling pathways stimulate different behaviorally relevant genes

Given our above observations, we explored which downstream neural events are elicited by Ara and Arb to achieve aggression and mating. First, to determine which brain nuclei expressing Ara and Arb are relevant to the display of these behaviors, we analyzed the expression of *fos* in each brain nucleus of males that mated with females and males that attacked other males, and compared the results with those of solitary males that neither mated nor attacked. In situ hybridization showed that *fos* expression was significantly higher in many brain regions of males that mated as compared with solitary males. These regions included the Vd/Vs/Vp in the ventral telencephalon, PMp, PPa, and PPp/PMm/PMg in the preoptic area, VM in the thalamus, and NAT, NVT, and NPT in the hypothalamus, most of which are components of the social behavior network (13, 14) (Fig. 5B and Fig. S8). For males that attacked other males, significantly higher *fos* expression was found in the Vd/Vs/Vp, PPa, NVT, and NPT (Fig. 5B and Fig. S8). These results suggest that one or more of these brain nuclei are sites of action of Ara and Arb signaling for aggression and mating, respectively.

Next, we explored which effector genes act downstream of Ara and Arb signaling in these brain nuclei. We focused on two neuropeptide genes, *vt* (encoding vasotocin) and *gal* (encoding galanin), which have been implicated in aggression and mating in many vertebrates, including medaka and other teleosts (19, 36, 37). In medaka, *vt* and *gal* are expressed in an androgen-dependent, and hence male-biased, manner in the posterior part of the NVT (pNVT) and PMp (pPMp), respectively (19, 21, 38), where increased *fos* expression was observed in our in situ hybridization analysis (Fig. 5B). Because *vt* neurons in the pNVT express *ara* but not *arb*, and *gal* neurons in the pPMp express *arb* but not *ara* (19, 21), we hypothesized that Ara signaling might facilitate male-directed aggression by inducing *vt* expression in the pNVT, and Arb signaling might facilitate mating with females by inducing *gal* expression in the pPMp.

We first tested these hypotheses by examining the expression of *vt* and *gal* in the brains of *ara*^−/−^ and *arb*^−/−^ males and females by in situ hybridization. The male-specific expression of *vt* in the pNVT was almost completely abolished in *ara*^−/−^ males (Fig. 5 C and D), but no significant differences between *ara*^+/+^ and *ara*^−/−^ males were observed in other brain nuclei (Fig. S9A). In contrast, the levels of *vt* expression in *arb*^−/−^ males were similar to those in *arb*^+/+^ males in the pNVT (Fig. 5 E and F), but slightly decreased in the PMp/PPa/PMm/PMg (Fig. S9 B and C). Expression of *gal* in the pPMp remained intact in *ara*^−/−^ males (Fig. 5 G and H), but was reduced in *arb*^−/−^ males to approximately one-third of the level in *arb*^+/+^ males (Fig. 5 I and J). In both *ara*^−/−^ males and *ara*^−/−^ females, reduced *gal* expression was instead observed in the aPMp/PPa and NAT/NVT/NRL (Fig. S9 D–F), while in *arb*^−/−^ males, no significant changes in *gal* expression were found outside the pPMp (Fig. S9G). These results demonstrate that the expression of *vt* in the pNVT and that of *gal* in the pPMp are critically and exclusively dependent on Ara and Arb signaling, respectively, and imply that the loss of this *vt* and *gal* expression might account for the behavioral deficits observed in *ara*^−/−^ and *arb*^−/−^ males, respectively.

Finally, we tested our hypothesis by administering Vt and Gal peptides to *ara*^−/−^ and *arb*^−/−^ males, respectively, to explore whether their behavioral deficits could be rescued. As predicted, Vt-treated *ara*^−/−^ males engaged in aggressive acts more frequently relative to non-treated controls, with significant differences for fin displays and circles (Fig. 5 K and L). By contrast, Gal treatment had no significant effect on any measure of mating or aggression in *arb*^−/−^ males (Fig. S10). Taken altogether, our results suggest that *vt* acts downstream of Ara signaling in the pNVT to mediate its stimulatory effects on male-directed aggression; however, it remains to be determined whether *gal* mediates the behavioral effects of Arb signaling.

## Discussion

Herein, we have explored the behavioral consequences of the loss of two AR subtypes in male medaka. Our results reveal, to our surprise, that males lacking Ara are less aggressive toward, and instead actively court, other males, while those lacking Arb are less motivated to mate with, and instead attack, females. These findings signify that, in male medaka, Ara- and Arb-mediated androgen signaling facilitate appropriate behavioral responses, while simultaneously suppressing inappropriate responses, to male and female conspecifics, respectively. It thus follows that the adaptive behavioral responses of males are shaped by the complementary action of two distinct androgen signaling pathways. Up until this study, mouse *trpc2*, which encodes a cation channel critical for signal transduction in the vomeronasal organ, has been the only known vertebrate gene that, when deleted, reliably causes males to court other males or attack females (39, 40). Male mice lacking *trpc2* are unable to discriminate between male and female conspecifics, indicating that this gene plays an essential role in sex discrimination (39, 40). In contrast, male medaka lacking *ara* or *arb* retain the ability to discriminate sex; therefore, these two genes most probably act in the circuitry controlling decision-making rather than in sex recognition.

Our results also suggest that Ara and Arb signaling inhibit each other to ensure mutually exclusive displays of mating and aggression. Considering that Ara and Arb suppress male-directed mating behavior and female-directed aggression, respectively, while facilitating aggression and mating, respectively, we postulate that Ara inhibits Arb signaling to prevent male–male mating, while Arb inhibits Ara signaling to prevent males attacking females. This reciprocal inhibition of Ara and Arb signaling probably involves interactions that occur between neurons, rather than within a single neuron, because Ara and Arb are expressed primarily in different neurons. This system evokes parallels with the reciprocal inhibition between *Esr1*-expressing neurons in the MPOA and VMHvl of male mice that functions in the decision to choose whether to mate or fight (5, 6). Such neuronal interactions involving sex steroid signaling may be conserved in vertebrates and represent a general mechanism by which males decide whether to mate or fight, although the specific sex steroid and receptor involved in this mechanism is like to vary among species. Given that both the PMp and NAT – the putative teleost homologs of the rodent MPOA and VMH, respectively (13, 41) – express *ara* and/or *arb*, the brain nucleus associated with this mechanism may be also conserved across species. This idea needs to be explored in future studies.

It may be relevant to note here that *Esr1*-expressing neurons in the mouse MPOA and VMHvl also express *Ar* (42). This raises the question of whether androgen/AR signaling may play a critical role in mediating male behavioral responses in mice as well as in medaka. In mice, however, the role of androgen/AR signaling in male behaviors is rather limited: unlike in many other vertebrates, it only increases the extent of behaviors (43); and indeed, male mice lacking *Ar* in the brain do not frequently court other males or attack females (44, 45). Instead, male mice lacking *Esr1* in inhibitory neuronal populations show aggression, albeit not significant, toward females (46). Therefore, it can be still concluded that the specific sex steroid signaling pathway playing a causal role in male behaviors varies among species. Furthermore, a recent study in male cichlid demonstrated that Ara is essential for mating, while either Ara or Arb is sufficient for aggression (24), suggesting that species differences may exist even among teleosts.

Another significant finding in this study is that the Ara and Arb signaling pathways are activated in females in response to exogenous 11KT administration, producing male-typical aggressive and mating behaviors, respectively. This finding suggests either that the functional neural circuits through which these signaling pathways direct male-typical behavioral responses during social encounters exist in the normal female brain, or that these circuits are rapidly organized by exogenous 11KT. Whichever is the case, these circuits must be one of the neural substrates through which changes in the sex steroid milieu of adult teleosts effectively reverse their sex-typical behaviors. Among vertebrates, teleosts are exceptionally sexually plastic in their behavior as adults; however, there is accumulating evidence to suggest that other species, including rodents, also have some degree of plasticity (e.g., 47–49). Further research in teleosts is likely to shed light on the neural basis of sexual differentiation and plasticity of sex-typical behaviors in vertebrates.

With the exception of some species such as rodents, androgen/AR signaling is essential for the expression of male-typical behaviors (15–18). Nevertheless, limited information is available on the downstream effectors that mediate the behavioral effects of androgen/AR signaling, including the direct targets of AR (50). Our current studies suggest that the neuropeptide Vt (vasotocin) acts downstream of Ara signaling in the pNVT to mediate its stimulatory effects on male-directed aggression. Importantly, *vt* is expressed exclusively in males in the pNVT and Ara can directly activate the transcription of *vt* (21); thus, it is highly likely that *vt* serves as a male-specific, direct target of Ara for male-typical aggression. The teleost NVT is considered homologous to the rodent anterior hypothalamus (AH), a known major site of action of VT relevant to male behaviors (13, 41). Studies in hamsters, for example, have shown that androgen administration increases the amount of VT peptide in the AH while concurrently increasing aggression, and that administration of a VT receptor antagonist to the AH diminishes androgen-induced aggression (51, 52). These observations in hamsters seem to be comparable to the present results in medaka. Hence, the neural mechanism whereby androgen/AR signaling in the pNVT/AH stimulates male-typical aggression via VT may be conserved across species.

We also showed that another neuropeptide, Gal (galanin), is a target of Arb signaling in the pPMp, but found no evidence that Gal mediates the behavioral effects of Arb signaling. In our previous study, loss of *gal* in medaka resulted in a marked reduction in male–male aggressive chases, suggesting that *gal* is involved in male aggression (19). However, the present results show that *arb*-deficient males engage normally in aggressive chases despite reduced *gal* expression in the pPMp. A recent study in midshipman fish (*Porichthys notatus*) suggests that Gal neurons in the pPMp play a role in male mating behavior (37), but our treatment of *arb*-deficient males with Gal did not restore their impaired mating behavior. One possible explanation for these discrepancies is that the reduction in *gal* expression in the pPMp of *arb*-deficient males (down to approximately one-third) was not sufficient to affect behavior. Further studies are needed to pinpoint the downstream effectors of Arb signaling.

In conclusion, our findings reveal that male medaka make adaptive decisions to mate or fight as a result of the activation of one of two functionally distinct androgen/AR signaling pathways, depending on the sex of the conspecific that they encounter. Furthermore, our findings suggest that these pathways inhibit each other to ensure mutually exclusive displays of mating and aggression. Although the relevance of androgen/AR signaling in vertebrate male behaviors has long been recognized, to our knowledge, this study is the first to show that androgen/AR signaling stimulates male-typical mating and aggression while simultaneously suppressing these opponent behaviors toward inappropriate targets. Last but not least, the findings that the deficiency of a single gene in medaka can cause male–male mating (in the case of *ara*; this study) or female–female mating (in the case of *esr2b*; 20) provide important insights into the neural substrates underlying sexual orientation and their evolutionary aspects.

## Materials and Methods

### Animals

Wild-type and *ara-*, *arb-*, and *esr2b*-deficient medaka (20) on the d-rR genetic background were raised at 28°C with a 14-hour light/10-hour dark photoperiod. They were fed with live *Artemia nauplii* and dry food (Otohime; Marubeni Nisshin Feed, Tokyo, Japan) 3–4 times a day. Sexually mature, spawning adults (aged 2–6 months) were used in all experiments and assigned randomly to experimental groups. The fish used in each experiment were age-matched, co-housed siblings to control for genetic and environmental confounding. Tissue sampling was consistently performed 1–3 hours after light onset. All experimental procedures involving animals were performed in accordance with the University of Tokyo Institutional Animal Care and Use Committee guidelines.

### Generation of gene-deficient medaka

*ara*- and *arb*-deficient medaka were generated by CRISPR/Cas9-mediated genome editing. Two and one CRISPR RNAs (crRNAs) were designed to target the predicted DNA-binding domains of *ara* and *arb* (22), respectively (Fig. S1A for *ara*; Fig. S2A for *arb*). The crRNAs and trans-activating crRNA (tracrRNA) were synthesized by Fasmac (Kanagawa, Japan) and injected along with Cas9 protein (Nippon Gene, Tokyo, Japan) into the cytoplasm of embryos at the one- or two-cell stage. Upon reaching adulthood, injected fish were outcrossed with wild-type fish, and the resulting progeny were tested for target site mutations by T7 endonuclease I assay (53), followed by direct sequencing. For *ara* targeting, two founders were identified that reproducibly produced progeny with deleterious mutations: one produced progeny carrying a 327-bp deletion/1-bp insertion (Δ326), which removed both the DNA- and ligand-binding domains; the other produced progeny carrying a 338-bp deletion/13-bp insertion (Δ325), which removed the N-terminal half of the DNA-binding domain (Fig. S1). Similarly, for *arb* targeting, two founders were identified that produced progeny carrying, respectively, 10-bp and 11-bp deletions (Δ10 and Δ11), both of which removed the DNA- and ligand-binding domains (Fig. S2). For both *ara* and *arb*, progeny from each founder were intercrossed to establish two independent mutant lines. Each line was maintained by intercrossing heterozygotes to obtain wild-type and homozygous siblings for experimental use. All fish were genotyped by PCR amplification of the target locus, followed by agarose gel electrophoresis (*ara*-deficient lines) or high-resolution melting analysis (*arb*-deficient lines) using the primers and probe listed in Table S2.

### Aggressive behavior test

Intrasexual aggressive behavior among grouped fish was assessed as described by Yamashita et al. (19). In brief, four fish of the same sex and genotype (*ara*- or *arb*-deficient fish or wild-type siblings), unfamiliar with one another, were placed together in a 2-liter rectangular tank 1 hour after light onset (2 hours after treatment in tests using Vt and Gal). After acclimation to the test tank for 1 min, fish were allowed to interact for 15 min. All interactions were recorded with a digital video camera (HC-V360MS, HC-VX985M, or HC-W870M; Panasonic, Tokyo, Japan), and the total number of each aggressive act (chase, fin display, circle, strike, and bite) displayed by the four fish in the tank was counted manually from the video recordings. In all video analyses, the researcher was blind to the fish genotype and treatment.

Aggressive behavior was also tested individually by pairing each fish with a stimulus fish. On the day before behavioral testing, a focal (*ara*- or *arb*-deficient fish or wild-type siblings) and stimulus (wild-type male in the tests shown in Fig. S4 A–F; wild-type female in Fig. 4 A–F, J, and K and Fig. S6 A–E; and *esr2b*-deficient female in Fig. S6 F–J) fish were placed in a 2-liter rectangular tank with a perforated transparent partition separating them. The exception was the test using 11KT-treated females, where each focal fish received 11KT continuously and was not introduced to the test tank until the day of testing to ensure the effectiveness of 11KT. The partition was removed 1 hour after light onset, and fish were allowed to interact for 15 min while their behavior was recorded as described above. The interaction time was increased to 30 min in the tests using stimulus females (wild-type or *esr2b*-deficient), because females were attacked relatively infrequently. The number of each aggressive act performed by the focal fish, and the percentage of focal fish exhibiting any aggressive act were calculated from the video recordings. In tests where the focal and stimulus fish were of the same sex, a small portion of the caudal fin of the stimulus fish was clipped to distinguish them.

Regarding the behavior of *arb*-deficient males toward wild-type stimulus females, the correlation between the number of aggressive acts during the interaction period and the number of wrapping attempts in the first 10 min of interaction, and the latency from the first mating act to the initiation of aggressive acts were further analyzed (note that the correlation analysis was not performed for *arb*-deficient males of the Δ11 line, because only 4 of 11 males showed wrappings). In these analyses, courtship displays and wrapping attempts, in addition to aggressive acts, were time-stamped in the first 15 min of interaction and used to create the raster plots shown in Fig. 4B.

### Mating behavior test

*ara*- and *arb*-deficient fish and wild-type siblings were tested for mating behavior toward a stimulus fish (wild-type female in the tests shown in Fig. 2 B–G, Fig. S4 J–M, and Fig. S10A; *esr2b*-deficient female in Fig. 2 H–J and Fig. S3; or wild-type male in Fig. 3 A–F and Fig. S5) using essentially the same procedure as described above for aggressive behavior, with the following exceptions. The partition was removed after 2 hours of treatment in the tests using Vt- and Gal-treated males. The interaction time was increased to 30 and 120 min in the tests using, respectively, *esr2b*-deficient females as the stimulating fish and 11KT-treated females as the focal fish because the 15-min interaction time was insufficient.

The latencies of the focal fish to initiate followings, courtship displays, and wrappings, and to spawn were calculated from video recordings. In tests using *esr2b*-deficient females and wild-type males (which were unreceptive to male courtship and did not spawn) as stimulus fish, the number of courtship displays and wrapping attempts refused by the stimulus fish was calculated instead of the latency to spawn. In tests where the focal and stimulus fish were both males, aggressive acts, in addition to courtship displays and wrapping attempts, were time-stamped in the first 5 min of interaction and used to create the raster plots shown in Fig. 3B. Note that the latency to initiate followings was not calculated in these tests because it was sometimes difficult to distinguish between followings for mating and aggressive chases.

### Drug treatment

Females of the *ara*-deficient (Δ326) and *arb*-deficient (Δ10) lines were treated with 100 ng/ml of 11KT (Denis Pharma, Tokyo, Japan) by immersion in water for 14–16 days and then tested for aggressive and mating behaviors as described above. In separate experiments, *ara*-deficient (Δ326) and *arb*-deficient (Δ10) males were treated intraperitoneally with 1 μl of an 8 μg/ml solution of synthetic medaka Vt and Gal peptides (Scrum, Tokyo, Japan), respectively, or with vehicle (phosphate-buffered saline) alone 1–1.5 hours after light onset. Aggressive and mating behaviors were evaluated 2 hours after treatment as described above.

### Three-chamber test

The test apparatus consisted of a rectangular tank (165 mm long by 190 mm wide, filled with water to 30 mm) divided into three chambers by two perforated transparent partitions positioned to give a central (test) chamber of 105 mm and two side chambers of 30 mm in length (Fig. 3 G and J and Fig. 4G). Focal males (*ara*- and *arb*-deficient males or wild-type siblings) were individually placed in the test chamber, while a wild-type stimulus female, unfamiliar to the focal males, was placed in one side chamber. The other side chamber either contained a wild-type unfamiliar male or was left empty. After acclimation for 1 min, the focal males were allowed to swim freely in the test chamber for 10 min while their behavior was recorded. All tests were conducted 1–3 hours after light onset. The position of the stimulus male and female was alternated every two trials to control for any side-preference bias. The location of the focal males was tracked from the video recordings and the time spent in each location was calculated using UMATracker (54). The resulting data were used to generate heat maps for visual analysis and to calculate the difference in time spent near the stimulus male versus the stimulus female.

### Analysis of *fos* expression

The focal fish (wild-type male) was placed alone or paired with a wild-type stimulus female or male in a 2-liter rectangular tank with a perforated transparent partition separating them. The partition was removed to allow fish to interact 0.5–1.5 hours after light onset on the following day. The brain of each paired focal male was sampled 30 min after it spawned with the stimulus female or exhibited any aggressive act toward the stimulus male, and analyzed for *fos* expression by in situ hybridization (see below). The brains of solitary males were sampled 30 min after removal of the partition and processed likewise.

### Single-label in situ hybridization

DNA fragments corresponding to nucleotides 16–1030 (1015 bp) of the *ara* cDNA (GenBank accession number: NM_001122911), 53–1233 (1181 bp) of the *arb* cDNA (NM_001104681), 20–1223 (1204 bp) of the *fos* cDNA (NM_001252234), 1–845 (845 bp) of the *vt* cDNA (NM_001278891), and 5–533 (529 bp) of the *gal* cDNA (LC532140) were PCR-amplified and transcribed in vitro to generate digoxigenin (DIG)-labeled cRNA probes using T7 RNA polymerase and DIG RNA Labeling Mix (Roche Diagnostics, Basel, Switzerland).

Single-label in situ hybridization was performed as described previously (28). In brief, brains dissected from males of the *ara*-deficient (Δ326) line (for analysis of *arb*, *vt*, and *gal*), *arb*-deficient (Δ10) line (*ara*, *vt*, and *gal*), and wild-type strain (*fos*) were fixed in 4% paraformaldehyde (PFA) and embedded in paraffin. Serial sections (10-μm thick) were cut in the coronal plane and hybridized with one of the DIG-labeled probes described above. Hybridized probes were detected using an alkaline phosphatase-conjugated anti-DIG antibody (RRID: AB_514497; Roche Diagnostics) with nitro blue tetrazolium/5-bromo-4-chloro-3-indolyl phosphate (NBT/BCIP) substrate (Roche Diagnostics). The color was allowed to develop for 1 hour (*gal*), 6 hours (*fos* and *vt*), or overnight (*ara* and *arb*). Brain nuclei were identified using medaka brain atlases (55, 56). Images were acquired with a virtual slide microscope (VS120; Olympus, Tokyo, Japan), and the total area of expression signal in each brain nucleus was calculated using Olyvia software (Olympus).

### Double-label in situ hybridization

The above-mentioned *ara* and *arb* cRNA probes were labeled, respectively, with fluorescein using T7 RNA polymerase and Fluorescein RNA Labeling Mix (Roche Diagnostics) and with DIG as described above.

Wild-type male brains were fixed in 4% PFA, embedded in paraffin, and coronally sectioned (10-μm thick). The sections were hybridized simultaneously with the fluorescein-labeled *ara* and DIG-labeled *arb* probes. Fluorescein was detected with a horseradish peroxidase-conjugated anti-fluorescein antibody (RRID: AB_2737388; PerkinElmer, Waltham, MA) and visualized by using the TSA Plus Fluorescein System (PerkinElmer); DIG was detected with a mouse anti-DIG antibody (RRID: AB_304362; Abcam, Cambridge, UK) and visualized using the Alexa Fluor 555 Tyramide SuperBoost Kit, goat anti-mouse IgG (Thermo Fisher Scientific, Waltham, MA, USA). Sections were counterstained with 4′,6-diamidino-2-phenylindole (DAPI) to identify cell nuclei. Fluorescent images were acquired with a confocal laser scanning microscope (Leica TCS SP8; Leica Microsystems, Wetzlar, Germany) with the following excitation/emission wavelengths: 405/410–480 nm (DAPI), 488/495–545 nm (fluorescein), and 552/620–700 nm (Alexa Fluor 555).

To calculate the percentage of neurons co-expressing *ara* and *arb*, at least 40 *ara*- and/or *arb*-expressing neurons from at least four different sections of each brain nucleus per fish were photographed and analyzed for overlapping expression of *ara* and *arb*.

### Statistical analysis

Data for continuous variables were expressed as mean ± standard error of the mean (SEM), with individual data points shown as dots. Categorical variables were expressed as percentages. Behavioral time-series data were analyzed using Kaplan-Meier plots with the inclusion of fish that did not exhibit the given act within the test period, in accordance with Jahn-Eimermacher et al. (57).

All statistics were analyzed using GraphPad Prism (GraphPad Software, San Diego, CA). Comparisons between two groups for continuous variables were performed using the unpaired two-tailed Student’s *t* test. Welch’s correction was applied if the F-test indicated that the variance between group was significantly different. Continuous variables in more than two groups were analyzed using one-way analysis of variance (ANOVA), followed by Dunnett’s post hoc test. If the Brown-Forsythe test indicated a significant difference in variance across groups, the data were instead analyzed using the non-parametric Kruskal-Wallis test followed by Dunn’s post hoc test. Differences between Kaplan-Meier curves were tested for significance using Gehan-Breslow-Wilcoxon test. A one-sample *t* test was used to test the null hypothesis that there would be no difference between the time spent on the stimulus female side and that spent on the male/empty side. The Pearson correlation coefficient was used to assess whether there was a linear correlation between the frequency of aggressive acts and that of wrapping attempts. Comparisons of categorical variables were performed using Fisher’s exact test. All data points were included in the analyses and no outliers were defined.

## Acknowledgments

We thank Dr. Yukiko Ogino for sharing information about the possible roles of *ara* and *arb* prior to publication; Dr. Hideaki Takeuchi for technical advice on three-chamber tests; and Akira Hirata, Kaoru Furukawa, and Tomiko Iba for assistance with medaka husbandry. This work was supported by the Ministry of Education, Culture, Sports, Science, and Technology (MEXT), Japan, and the Japan Society for the Promotion of Science (JSPS) (MEXT/JSPS grant numbers 21J20634 (to YN), 17H06429, and 23H02305 (to KO)).

## Notes

### Competing Interest Statement

The authors have declared no competing interest.

