## Supplementary Materials for "The decision of male medaka to mate or fight depends on two complementary androgen signaling pathways"

**Table S1. Abbreviations of brain nuclei.**

| Abbreviation | Full name | Location |
|---|---|---|
| aNVT | Anterior part of the NVT | Hypothalamus |
| aPMp | Anterior part of the PMp | Preoptic area |
| NAT | Anterior tuberal nucleus | Hypothalamus |
| NPT | Posterior tuberal nucleus | Hypothalamus |
| NRL | Lateral recess nucleus | Hypothalamus |
| NVT | Ventral tuberal nucleus | Hypothalamus |
| PMg | Gigantocellular portion of the magnocellular preoptic nucleus | Preoptic area |
| PMm | Magnocellular portion of the magnocellular preoptic nucleus | Preoptic area |
| PMp | Parvocellular portion of the magnocellular preoptic nucleus | Preoptic area |
| pNVT | Posterior part of the NVT | Hypothalamus |
| PPa | Anterior parvocellular preoptic nucleus | Preoptic area |
| pPMp | Posterior part of the PMp | Preoptic area |
| PPp | Posterior parvocellular preoptic nucleus | Preoptic area |
| SC | Suprachiasmatic nucleus | Preoptic area |
| VM | Ventromedial nucleus | Thalamus |
| Vp | Posterior nucleus of the ventral telencephalic area | Ventral telencephalon |
| Vs | Supracommissural nucleus of the ventral telencephalic area | Ventral telencephalon |
| Vv | Ventral nucleus of the ventral telencephalic area | Ventral telencephalon |

**Table S2. Primers and a probe used for genotyping of the *ara* and *arb* alleles.**

| Primer/probe | Allele | Direction | Purpose | Sequence (5′ to 3′) |
|---|---|---|---|---|
| Primer | *ara* | Forward | gDNA PCR | TGCTCGGGATGGAAGAAGATGT |
| Primer | *ara* | Reverse | gDNA PCR | TGCATTGAAGCGCAAACAGCTC |
| Primer | *ara* | Forward | CS | GGTTTCCAAAGAGGATGGACAG |
| Primer | *arb* | Forward | gDNA PCR | CCCAGTTTCAAGGCTTTAAGCC |
| Primer | *arb* | Reverse | gDNA PCR | AACTTCATGGGCAGTGTGATGC |
| Primer | *arb* | Forward | CS | TCCATAAGACACAGTCACTCCG |
| Primer | *arb* | Forward | HRM | TTCTTCATGTGTTTGACAGCAGGT |
| Primer | *arb* | Reverse | HRM | CCTTGCAACTTCCACAAGTTAGG |
| Probe | *arb* | Forward | HRM | CCCCCCTCAAAGGACGTGTCTGATCTGTTCAG |

gDNA PCR, PCR on genomic DNA; CS, cycle sequence; HRM, high-resolution melting analysis.

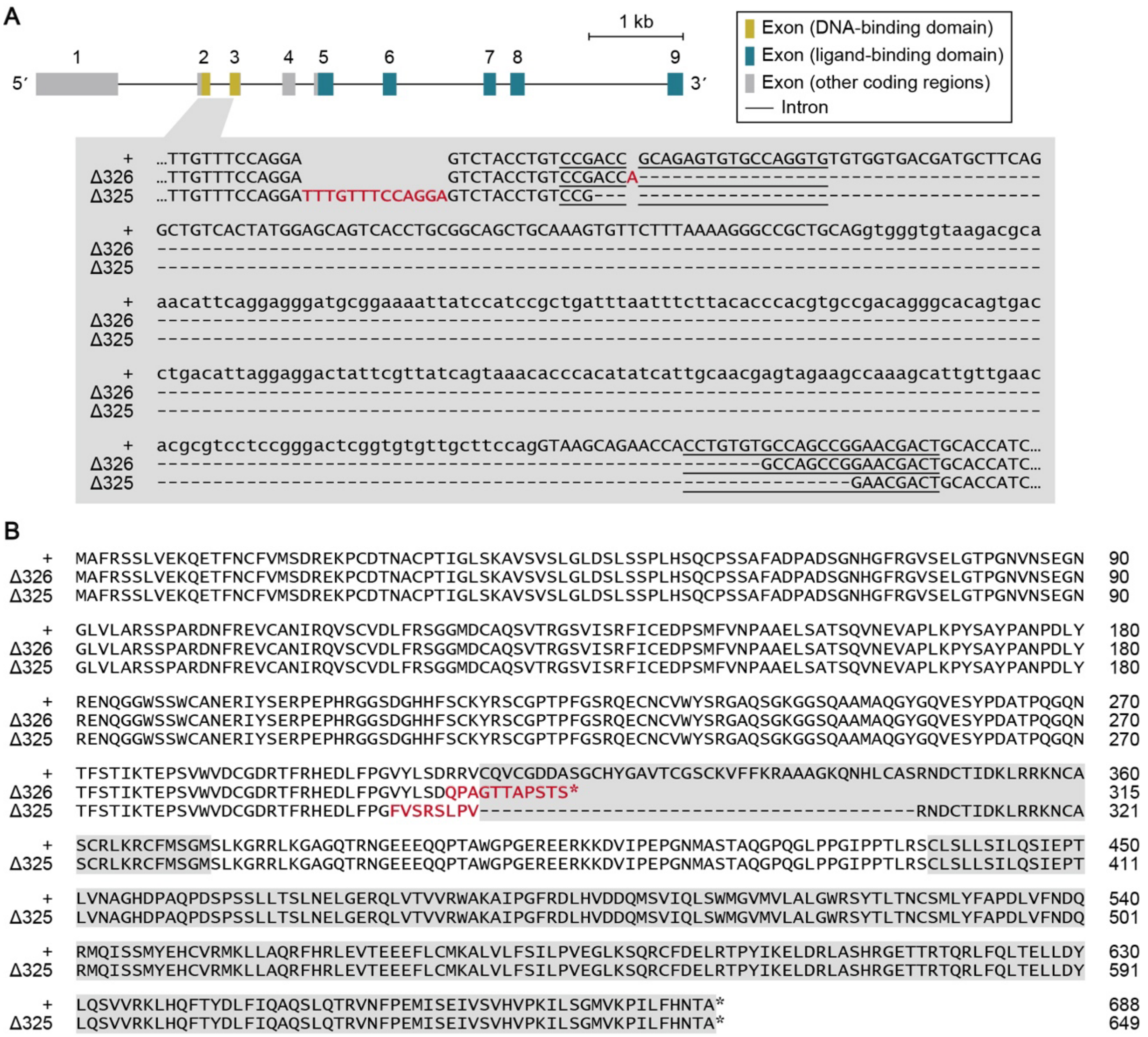


**Fig. S1 Generation of *ara*-deficient medaka.** (**A**) Schematic of the *ara* locus showing the CRISPR RNA target sites, which are enlarged to show the nucleotide sequences of the wild-type (+) and targeted (Δ326 and Δ325) alleles. Each exon is numbered above. Exon and intron sequences are denoted by capital and small letters, respectively. The CRISPR RNA target sequences are underlined. Inserted bases are indicated in red, and deleted bases are represented by dashes. (**B**) Amino acid alignment of the deduced wild-type (+) and targeted (Δ326 and Δ325) Ara proteins. The DNA-binding (residues 307–372 of wild-type Ara) and ligand-binding (residues 437–687 of wild-type Ara) domains are highlighted in gray. Residues generated as a consequence of frameshift are indicated in red. Asterisks denote stop codons.

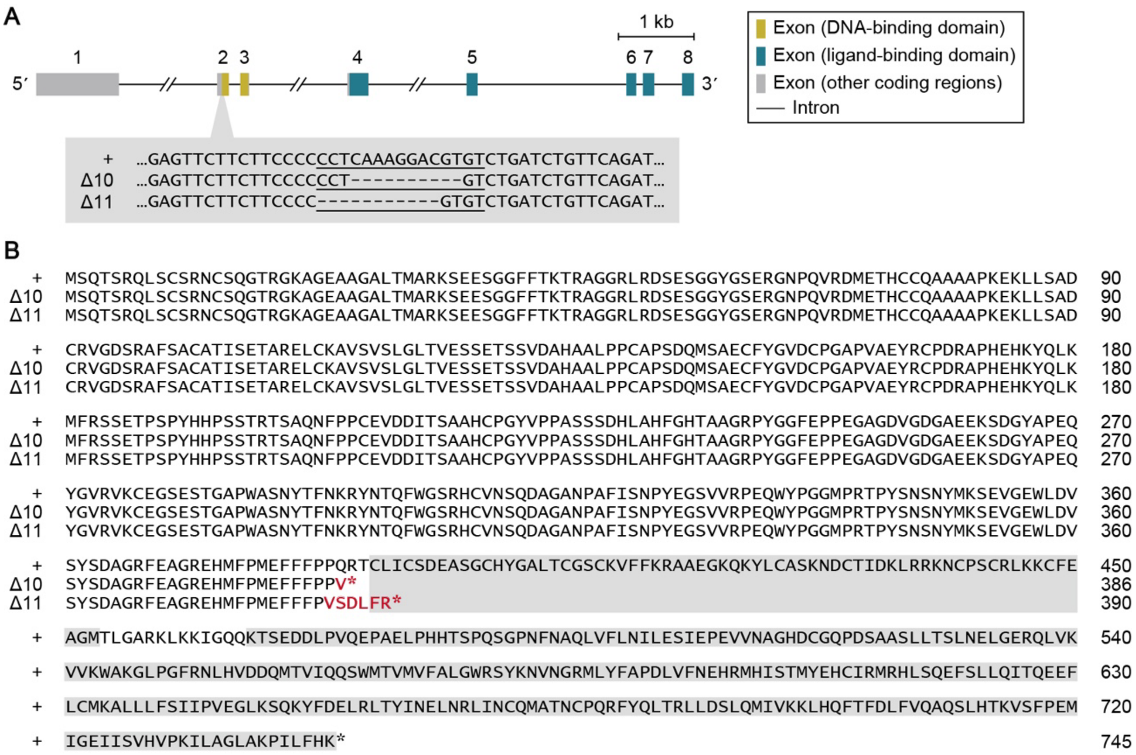


**Fig. S2 Generation of *arb*-deficient medaka.** (**A**) Schematic of the *arb* locus showing the CRISPR RNA target site, which is enlarged to show the nucleotide sequences of the wild-type (+) and targeted (Δ10 and Δ11) alleles. Each exon is numbered above. Exon and intron sequences are denoted by capital and small letters, respectively. The CRISPR RNA target sequence is underlined, and deleted bases are represented by dashes. (**B**) Amino acid alignment of the deduced wild-type (+) and targeted (Δ10 and Δ11) Arb proteins. The DNA-binding (residues 388–453 of wild-type Arb) and ligand-binding (residues 467–744 of wild-type Arb) domains are highlighted in gray. Residues generated as a consequence of frameshift are indicated in red. Asterisks denote stop codons.

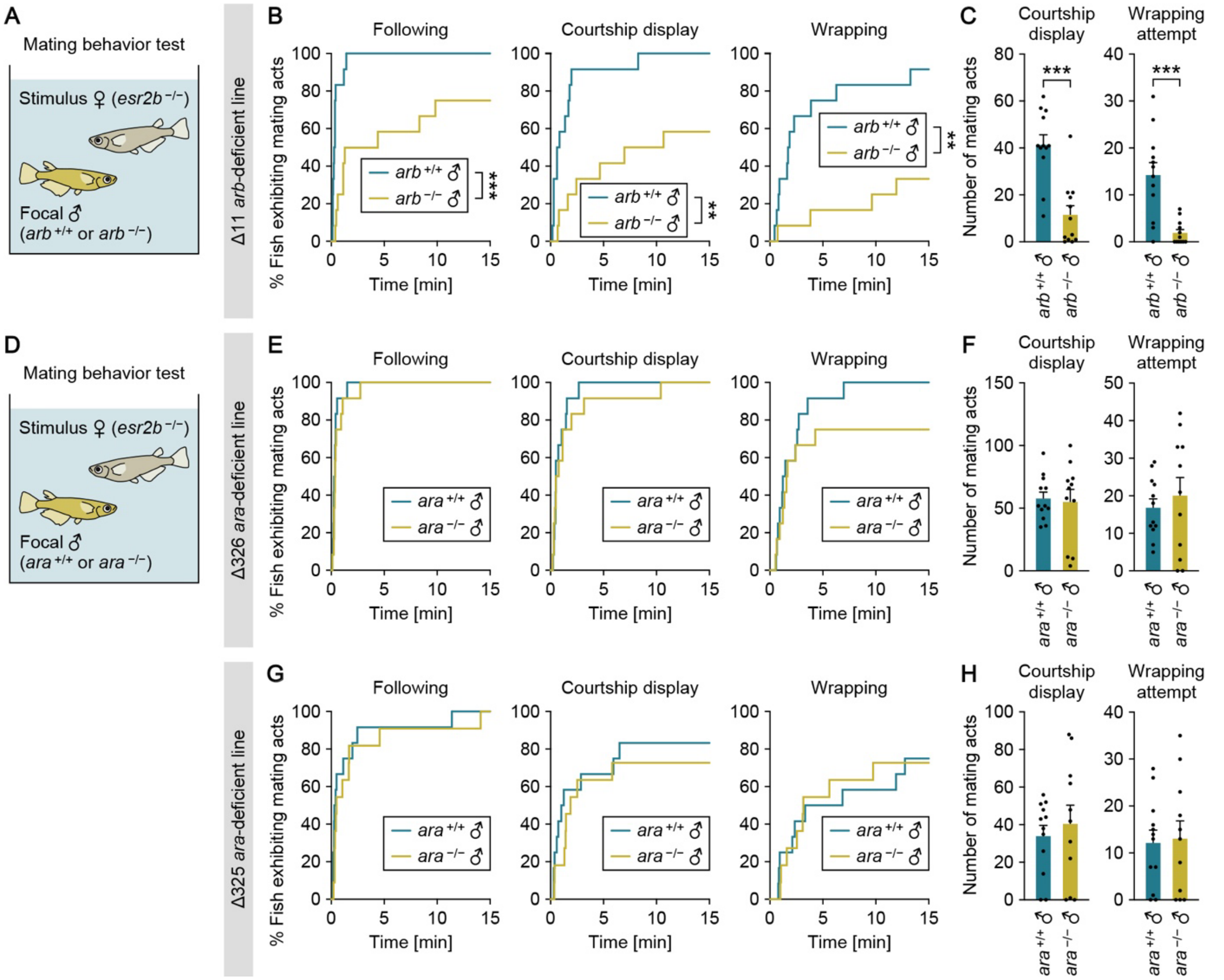


**Fig. S3 Mating behavior of *ara*- and *arb*-deficient males toward *esr2b*-deficient females.** (**A**) Set-up for testing the mating behavior of $arb^{+/+}$ and $arb^{-/-}$ males using an *esr2b*-deficient female as the stimulus. (**B**) Latency of the focal males (Δ11 line; n = 12 per genotype) to initiate each mating act toward the stimulus *esr2b*-deficient female. (**C**) Number of each mating act performed. (**D**) Set-up for testing the mating behavior of $ara^{+/+}$ and $ara^{-/-}$ males using an *esr2b*-deficient female as the stimulus. (**E**) Latency of the focal males (Δ326 line; n = 12 per genotype) to initiate each mating act toward the stimulus *esr2b*-deficient female. (**F**) Number of each mating act performed. (**G**) Latency of the focal males (Δ325 line; n = 12 and 11 for $arb^{+/+}$ and $arb^{-/-}$, respectively) to initiate each mating act toward the stimulus *esr2b*-deficient female. (**H**) Number of each mating act performed. Statistical differences were calculated by Gehan-Breslow-Wilcoxon test (B, E, and G), and unpaired *t* test, with Welch's correction where appropriate (C, F, and H). Error bars represent SEM. $^{**}P < 0.01$, $^{***}P < 0.001$.

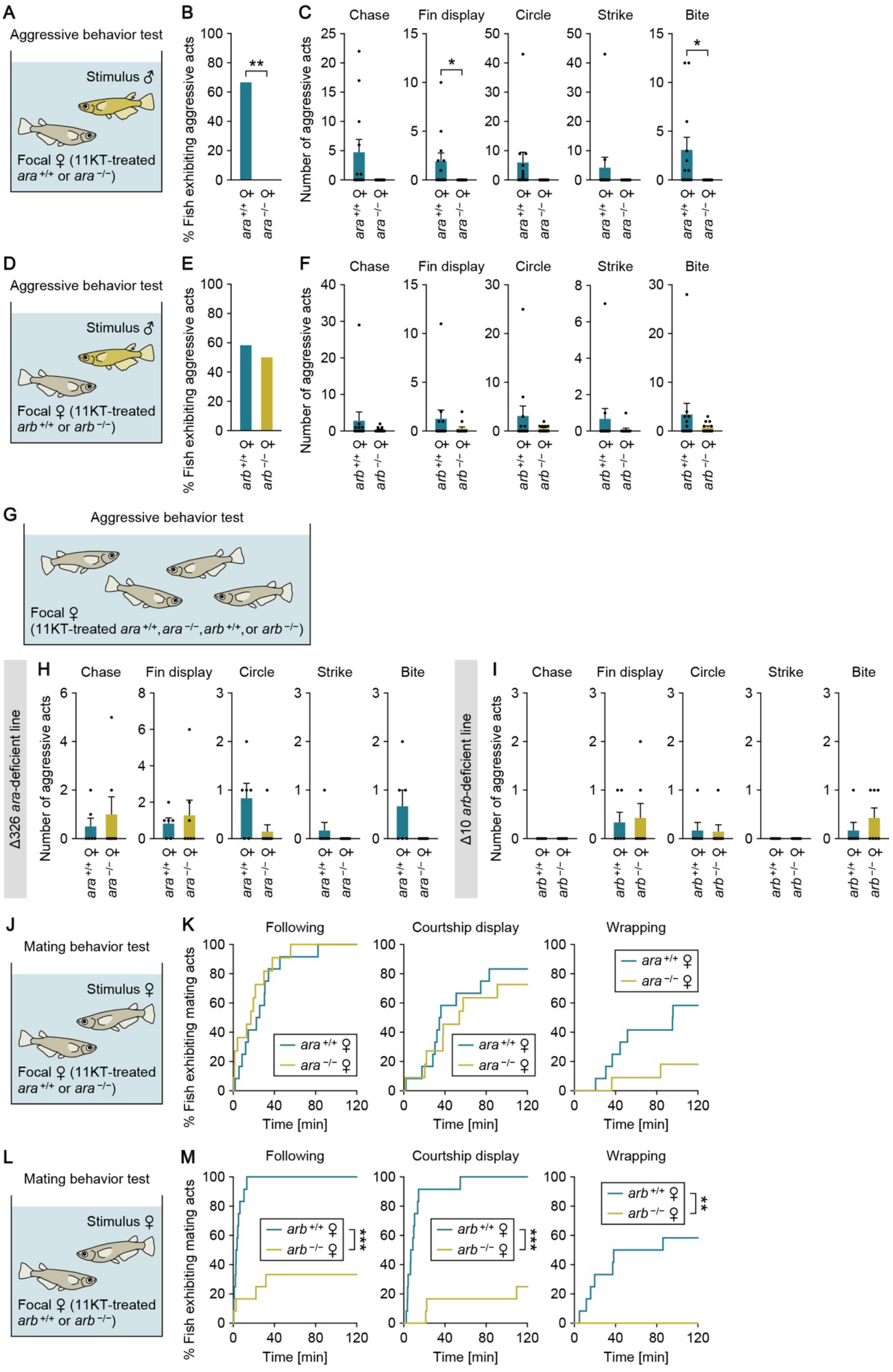
A
Aggressive behavior test
Stimulus ♂
Focal ♀ (11KT-treated ara+/+ or ara−/−)
B
% Fish exhibiting aggressive acts
**
ara+/+ ♀
ara−/− ♀
C
Chase
Fin display
Circle
Strike
Bite
Number of aggressive acts
*
D
Aggressive behavior test
Stimulus ♂
Focal ♀ (11KT-treated arb+/+ or arb−/−)
E
% Fish exhibiting aggressive acts
arb+/+ ♀
arb−/− ♀
F
Chase
Fin display
Circle
Strike
Bite
Number of aggressive acts
G
Aggressive behavior test
Focal ♀
(11KT-treated ara+/+, ara−/−, arb+/+, or arb−/−)
H
Δ326 ara-deficient line
Chase
Fin display
Circle
Strike
Bite
Number of aggressive acts
I
Δ10 arb-deficient line
Chase
Fin display
Circle
Strike
Bite
Number of aggressive acts
J
Mating behavior test
Stimulus ♀
Focal ♀ (11KT-treated ara+/+ or ara−/−)
K
Following
Courtship display
Wrapping
% Fish exhibiting mating acts
Time [min]
ara+/+ ♀
ara−/− ♀
L
Mating behavior test
Stimulus ♀
Focal ♀ (11KT-treated arb+/+ or arb−/−)
M
Following
Courtship display
Wrapping
% Fish exhibiting mating acts
Time [min]
arb+/+ ♀
arb−/− ♀
***
**

**Fig. S4 Adult androgen/AR signaling elicits male-typical behavioral responses even in females.** (**A**) Set-up for testing the aggressive behavior of 11KT-treated *ara*$^{+/+}$ and *ara*$^{-/-}$ females toward males. (**B**) Percentage of *ara*$^{+/+}$ and *ara*$^{-/-}$ females (Δ326 line; n = 12 per genotype) exhibiting any aggressive act toward the stimulus male. (**C**) Number of each aggressive act performed. (**D**) Set-up for testing the aggressive behavior of 11KT-treated *arb*$^{+/+}$ and *arb*$^{-/-}$ females toward males. (**E**) Percentage of *arb*$^{+/+}$ and *arb*$^{-/-}$ females (Δ10 line; n = 12 per genotype) exhibiting any aggressive act toward the stimulus male. (**F**) Number of each aggressive act performed. (**G**) Set-up for testing aggressive behavior among grouped females treated with 11KT. (**H**) Total number of each aggressive act observed among 11KT-treated *ara*$^{+/+}$ females and among 11KT-treated *ara*$^{-/-}$ females in the tank (Δ326 line; n = 6 and 7 for *ara*$^{+/+}$ and *ara*$^{-/-}$, respectively). (**I**) Total number of each aggressive act observed among 11KT-treated *arb*$^{+/+}$ females and among 11KT-treated *arb*$^{-/-}$ females in the tank (Δ10 line; n = 6 and 7 for *arb*$^{+/+}$ and *arb*$^{-/-}$, respectively). (**J**) Set-up for testing the mating behavior of 11KT-treated *ara*$^{+/+}$ and *ara*$^{-/-}$ females toward other females. (**K**) Latency of the focal females (Δ326 line, n = 12 and 11 for *ara*$^{+/+}$ and *ara*$^{-/-}$, respectively) to initiate each mating act toward the stimulus female. (**L**) Set-up for testing the mating behavior of 11KT-treated *arb*$^{+/+}$ and *arb*$^{-/-}$ females toward other females. (**M**) Latency of the focal females (Δ10 line; n = 12 per genotype) to initiate each mating act toward the stimulus female. Statistical differences were calculated by Fisher's exact test (B and E), unpaired *t* test, with Welch's correction where appropriate (C, F, H, and I), and Gehan-Breslow-Wilcoxon test (K and M). Error bars represent SEM. $*P < 0.05$, $**P < 0.01$, $***P < 0.001$.

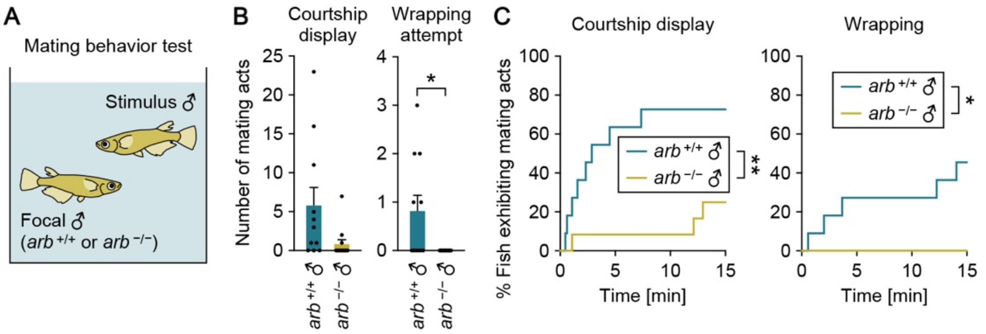


**Fig. S5 Mating behavior of *arb*-deficient males toward other males.** (**A**) Set-up for testing the mating behavior of $arb^{+/+}$ and $arb^{-/-}$ males toward other males. (**B**) Number of each mating act performed by the focal males (Δ10 line; n = 11 and 12 for $arb^{+/+}$ and $arb^{-/-}$, respectively). (**C**) Their latency to initiate each mating act.

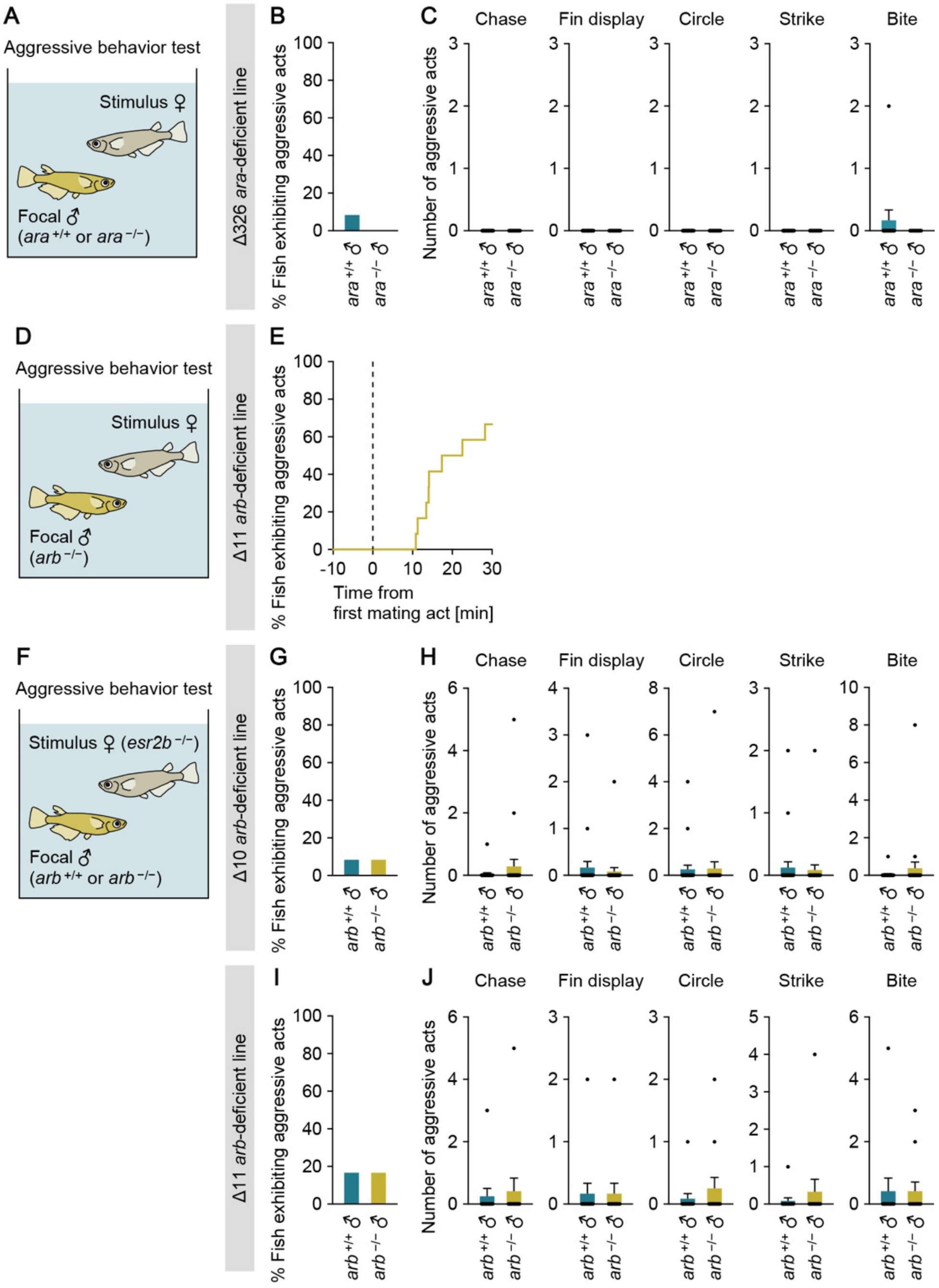


**Fig. S6 Aggressive behavior of *ara*- and *arb*-deficient males toward females.** (**A**) Set-up for testing the aggressive behavior of $ara^{+/+}$ and $ara^{-/-}$ males toward females. (**B**) Percentage of $ara^{+/+}$ and $ara^{-/-}$ males (Δ326 line; n = 12 per genotype) exhibiting any aggressive act toward the stimulus female. (**C**) Number of each aggressive act performed. (**D**) Set-up for testing the aggressive behavior of $arb^{-/-}$ males toward females. (**E**) Latency of $arb^{-/-}$ males (Δ11 line; n = 12) from the first mating act to initiate aggressive acts. (**F**) Set-up for the additional aggressive behavior test using an *esr2b*-deficient female as the stimulus.

(**G**) Percentage of *arb*$^{+/+}$ and *arb*$^{-/-}$ males (Δ10 line; n = 24 per genotype) exhibiting any aggressive act toward the stimulus *esr2b*-deficient female. (**H**) Number of each aggressive act performed. (**I**) Percentage of *arb*$^{+/+}$ and *arb*$^{-/-}$ males (Δ11 line; n = 12 per genotype) exhibiting any aggressive act toward the stimulus *esr2b*-deficient female. (**J**) Number of each aggressive act performed. Statistical differences were calculated by Fisher's exact test (B, G, and I) and unpaired *t* test, with Welch's correction where appropriate (C, H, and J). Error bars represent SEM.

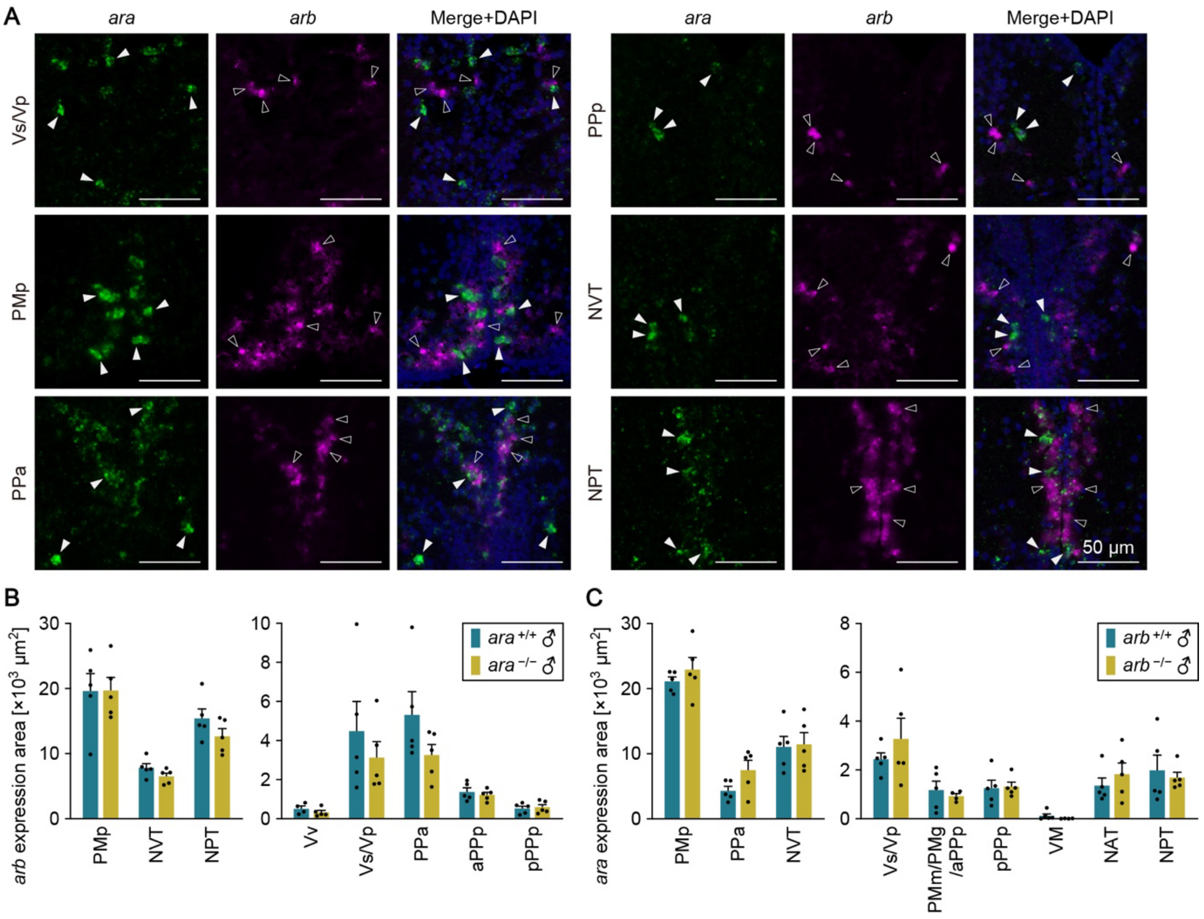


**Fig. S7 Independent association between *ara* and *arb* expression.** (**A**) Representative images of the expression of *ara* and *arb* in each brain nucleus. In each row, left panels show *ara* expression (green), middle panels show *arb* expression (magenta), and right panels show the merged images with DAPI staining (blue). White arrowheads indicate neurons expressing *ara* only; black arrowheads indicate neurons expressing *arb* only. All scale bars are 50 µm. (**B**) Total area of *arb* expression signal in each brain nuclei of $ara^{+/+}$ and $ara^{-/-}$ males (Δ326 line; n = 5 except for the Vv of $ara^{+/+}$ males, where n = 4). (**C**) Total area of *ara* expression signal in each brain nuclei of $arb^{+/+}$ and $arb^{-/-}$ males (Δ10 line; n = 5 except for the PMm/PMg/aPPp and VM of $arb^{-/-}$ males, where n = 4). For abbreviations of brain nuclei, see Table S1. Statistical differences were calculated by unpaired *t* test, with Welch's correction where appropriate (B and C). Error bars represent SEM.

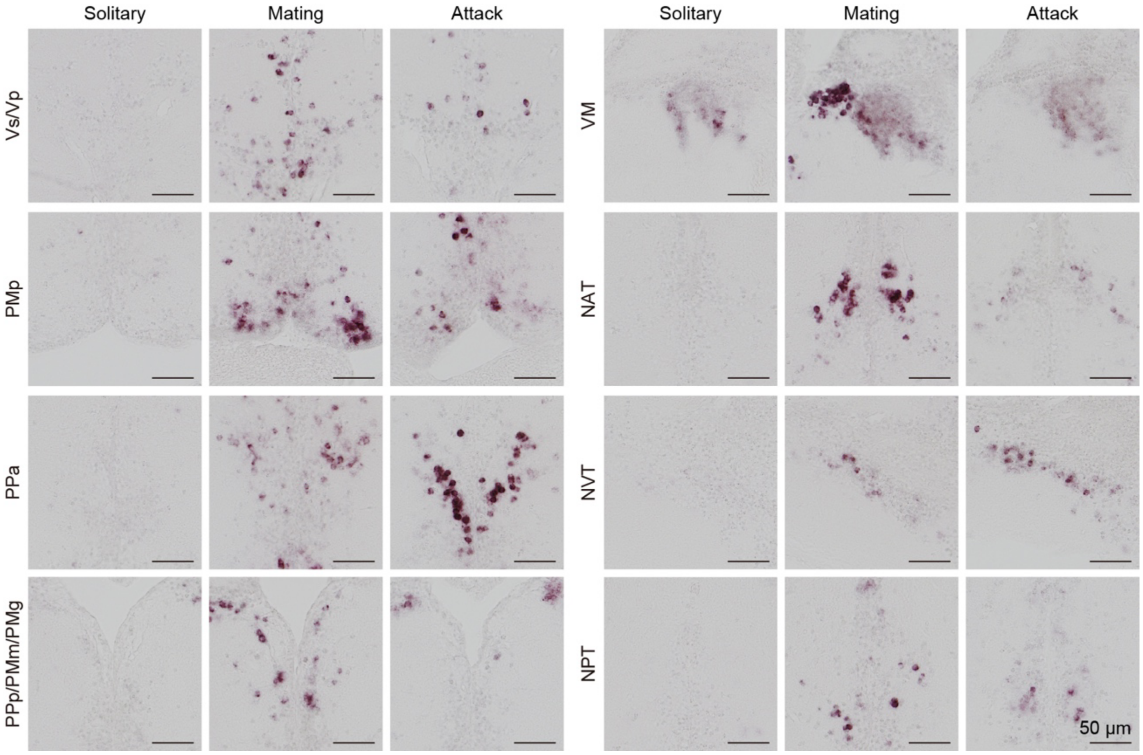


**Fig. S8 Increased *fos* expression in each brain nuclei upon mating and attack.** Representative images of *fos* expression in brain nuclei (where *ara* and/or *arb* are expressed) of wild-type solitary males, males that mated with females, and males that attacked other males. All scale bars are 50 μm. For abbreviations of brain nuclei, see Table S1.

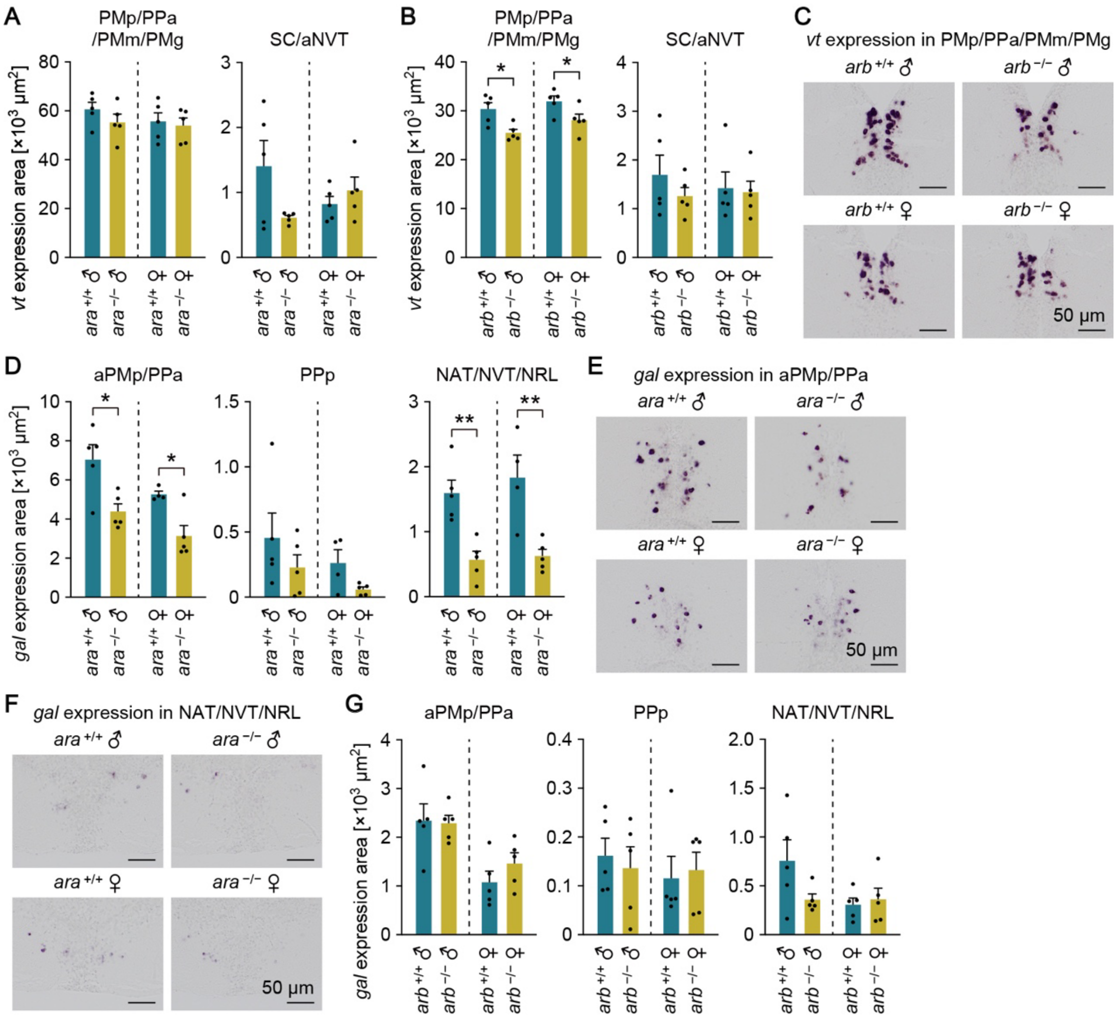


**Fig. S9 Expression of *vt* and *gal* in the brain of *ara*- and *arb*-deficient medaka.** (**A**) Total area of *vt* expression signal in the PMp/PPa/PMm/PMg and SC/aNVT of males and females of $ara^{+/+}$ and $ara^{-/-}$ fish (Δ326 line; n = 5 per sex per genotype). (**B**) Total area of *vt* expression signal in the PMp/PPa/PMm/PMg and SC/aNVT of males and females of $arb^{+/+}$ and $arb^{-/-}$ fish (Δ10 line; n = 5 per sex per genotype). (**C**) Representative images of *vt* expression in the PMp/PPa/PMm/PMg of these fish. (**D**) Total area of *gal* expression signal in the aPMp/PPa, PPp, and NAT/NVT/NRL of males and females of $ara^{+/+}$ and $ara^{-/-}$ fish (Δ326 line; n = 5 per sex per genotype, except n = 4 for $ara^{+/+}$ females). (**E** and **F**) Representative images of *gal* expression in the aPMp/PPa (E) and NAT/NVT/NRL (F) of these fish. (**G**) Total area of *gal* expression signal in the aPMp/PPa, PPp, and NAT/NVT/NRL of males and females of $arb^{+/+}$ and $arb^{-/-}$ fish (Δ10 line; n = 5 per sex per genotype). All scale bars are 50 μm. For abbreviations of brain nuclei, see Table S1. Statistical differences were calculated by unpaired *t* test, with Welch's correction where appropriate (A, B, D, and G). Error bars represent SEM. **P* < 0.05, ***P* < 0.01.

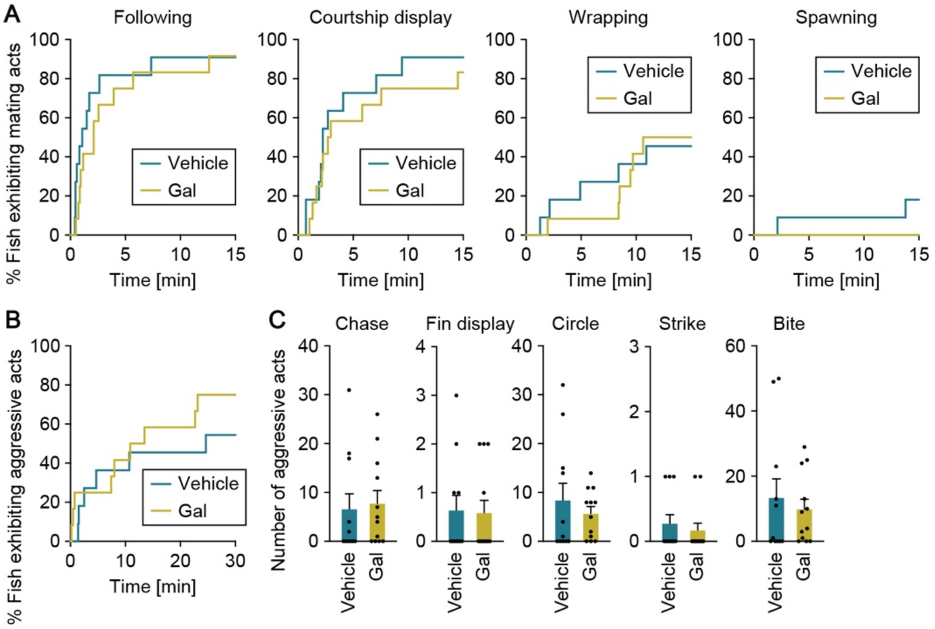


**Fig. S10 Mating and aggressive behaviors of Gal-treated *ara*-deficient males toward females.** (**A** and **B**) Latency of *arb*$^{-/-}$ males (Δ10 line) treated with vehicle alone (n = 11) or Vt peptide (n = 12) to initiate each mating act (A) and aggressive acts (B) toward the stimulus female. (**C**) Number of each aggressive act performed. Statistical differences were calculated by Gehan-Breslow-Wilcoxon test (A and B) and unpaired *t* test, with Welch's correction where appropriate (C). Error bars represent SEM.
